# Human iPSC-based coculture model reveals neuroinflammatory crosstalk between microglia and astrocytes

**DOI:** 10.1101/2025.01.03.628608

**Authors:** Iisa Tujula, Tanja Hyvärinen, Johanna Lotila, Julia Rogal, Dimitrios Voulgaris, Lassi Sukki, Kaisa Tornberg, Katri Korpela, Henna Jäntti, Tarja Malm, Anna Herland, Pasi Kallio, Susanna Narkilahti, Sanna Hagman

**Affiliations:** Neuroimmunology research group, Faculty of Medicine and Health Technology, Tampere University, Tampere, Finland; Science for Life Laboratory, Division of Nanobiotechnology, Department of Protein Science, Royal Institute of Technology (KTH), 171 65, Solna, Sweden; AIMES - Center for the Advancement of Integrated Medical and Engineering Sciences at Karolinska Institutet and KTH Royal Institute of Technology, Stockholm, Sweden; Department of Neuroscience, Karolinska Institutet, SE-171 77, Stockholm, Sweden; Micro and Nanosystems Research Group, Faculty of Medicine and Health Technology, Tampere University, Tampere, Finland; Neuroinflammation research group, Faculty of Health Sciences, A.I. Virtanen Institute for Molecular Sciences, University of Eastern Finland, Kuopio, Finland; NeuroGroup, Faculty of Medicine and Health Technology, Tampere University, Tampere, Finland

**Keywords:** Astrocytes, Microglia, iPSC, Disease modeling, Neuroinflammation, Glial crosstalk, Microfluidic chip platform, Coculture

## Abstract

**Background:** Microglia and astrocytes have been implicated as central mediators of neuroinflammatory processes in several neurodegenerative diseases. However, their intricate crosstalk and contributions to pathogenesis remain elusive, highlighting the need for innovative *in vitro* approaches for investigating glial interactions in neuroinflammation. The aim of this study was to develop advanced human-based glial coculture models to explore the inflammatory roles and interactions of microglia and astrocytes *in vitro*.

**Methods:** We utilized human induced pluripotent stem cell (iPSC)-derived microglia and astrocytes cultured both in conventional culture dishes and in a compartmentalized microfluidic chip coculture platform. This novel platform features separate compartments for both cell types, enabling the creation of fluidically isolated microenvironments with spontaneous migration of microglia toward astrocytes through interconnecting microtunnels. To induce inflammatory activation, glial cultures were stimulated with lipopolysaccharide (LPS), a combination of tumor necrosis factor-α (TNF-α) and interleukin-1β (IL-1β), or interferon-γ (IFN-γ) for 24 hours. The glial activation and crosstalk were analyzed with immunocytochemistry, the secretion of inflammatory factors from the culture media was measured, and microglial migration was quantified.

**Results:** Microglia–astrocyte cocultures were successfully generated in both conventional cultures and the microfluidic chip platform. Inflammatory stimulation with LPS and TNF-α/IL-1β elicited cell type-specific responses in microglia and astrocytes, respectively. Notably, the levels of secreted inflammatory mediators were altered under coculture conditions, revealing significant glial crosstalk. Utilization of our microfluidic coculture platform facilitated the study of microglial migration and glial activation within distinct inflammatory microenvironments. Microglia migrated efficiently toward the astrocyte compartment, and the chemoattractant adenosine diphosphate (ADP) notably increased microglial migration within this platform. Furthermore, inflammatory stimulation of the microfluidic chip cocultures successfully recapitulated glial crosstalk, revealing unique responses. This crosstalk was associated with elevated levels of complement component C3 in the cocultures, emphasizing the intricate interplay between microglia and astrocytes under inflammatory conditions.

**Conclusions:** Our results depict an elaborate molecular crosstalk between inflammatory microglia and astrocytes, providing evidence of how glial cells orchestrate responses during neuroinflammation. Importantly, we demonstrate that the microfluidic coculture platform developed in this study for microglia and astrocytes provides a more functional and enhanced setup for investigating inflammatory glial interactions *in vitro*.

## Background

Glial cells are increasingly recognized as not only passive support cells but also active mediators of neuroinflammation in various neurodegenerative diseases, including Alzheimer’s disease (AD), Parkinson’s disease (PD), amyotrophic lateral sclerosis (ALS) and multiple sclerosis (MS)^1,2^. Though these diseases occur via distinct pathogenetic mechanisms, such as protein aggregation, autoimmune reactions or genetic mutations, they all exhibit chronic neuroinflammation as a common feature. The key cell types that mediate neuroinflammatory reactions are microglia and astrocytes, the glial cells of the central nervous system (CNS). The glial cells play pivotal roles in maintaining CNS homeostasis, supporting neuronal function, and mediating inflammatory responses^3^. Upon CNS insult, microglia and astrocytes may adopt a diverse range of cellular states, resulting in morphological, molecular, and functional changes in response to pathological environmental cues^4,5^. The context-specific molecular mechanisms that drive inflammatory activation and dynamic neuroimmune responses of microglia and astrocytes are currently the focus of extensive research^6^.

Microglia and astrocytes do not function independently but modulate the functions of the other through tight bidirectional communication by soluble factors such as cytokines, chemokines, and growth factors^7,8^. As primary immune cells of the CNS, microglia are highly motile, constantly monitoring their environment and are the initial responders to CNS insults^7^. While many studies place microglia as the primary drivers of neuroimmune responses, evidence suggests that astrocytes can also serve as a significant source of cytokines and chemoattractants. As a result, reactive glial cells may lose their ability to perform homeostatic functions and gain neuroprotective or detrimental roles in the pathological environment^7,8^. The importance of bidirectional communication between astrocytes and microglia has been elegantly demonstrated in a previous study^9^, in which a specific reactive astrocyte state was induced by the release of cytokines, including tumor necrosis factor (TNF)-α, interleukin (IL)-1α and the complement component C1q, from microglia. These cytokines have been shown to upregulate complement component 3 (C3), contributing to neuronal death and synapse loss across different neurodegenerative disorders^9^. However, the underlying mechanisms of this elaborate inflammatory crosstalk between glial cells and its contribution to the spatiotemporal progression of neuroinflammatory pathologies remain unclear.

Recent breakthroughs in human induced pluripotent stem cell (iPSC) differentiation techniques have enabled the *in vitro* study of human microglia^10–12^ and astrocytes^13–15^, providing valuable insights into the pathogenic mechanisms of neurodegenerative diseases. The neuroinflammatory functions of iPSC-derived glial cells have been studied with approaches ranging from conventional monoculture- and coculture models to the use of conditioned media or more complex 3D neural organoids^6^. Similarly, a range of stimuli, including cytokines, aggregated proteins and endotoxins, have been used to initiate specific inflammatory responses in microglia and astrocytes. The power of disease-specific iPSC models in understanding neuroinflammation was demonstrated in our recent study, in which we showed that iPSC– microglia derived from patients with MS exhibit cell–intrinsic immune activation state evidenced by transcriptional changes and altered cytokine release, which potentially contribute to the neuroinflammatory environment in MS^16^. Moreover, some studies have also been performed using more complex cell culture setups, such as the triculture of neurons, astrocytes and microglia, to investigate neuroinflammation in AD. However, as the complexity of coculture models increases, so does the difficulty of identifying cell type-specific responses and certain cell–cell interactions^6^.

Conventional culture dishes offer limited control over culture design and cannot replicate the distinct microenvironments of the CNS^17^. Microfluidic technology has enabled the production of compartmentalized microscale culture platforms that direct and confine cell growth, allowing for the study of cell–cell interactions^17^. To overcome these limitations, we and others have utilized microfluidic chips for various CNS applications, including the isolation of neuronal axons and networks^18,19 20^, the creation of pathologically relevant disease models, such as for AD and PD, by studying the effects of aggregated proteins^21–23^, and the investigation of interactions between inflammatory glia and neuronal cells^24,25^. In our previous study, we demonstrated the effects of reactive astrocytes and the neuroinflammatory environment on axonal growth using a microfluidic chip platform^26^. Therefore, there is an urgent need to develop completely human-based neuroinflammatory models that can elucidate the cellular mechanisms involved and replicate the intricate interactions of glial cells.

In the present study, our aim was to investigate the inflammatory responses and crosstalk of cocultured iPSC-derived microglia and astrocytes. In addition to studying the effects of a panel of inflammatory stimulants in conventional cultures, we developed a novel microfluidic platform with astrocytes and microglia. The compartmentalized design enabled the establishment of separate, interconnected inflammatory environments with spontaneous microglial migration into the astrocyte compartment. The recruitment of microglia into activated astrocytes resulted in modulated immune responses that effectively recapitulated and, in some cases, enhanced glial crosstalk in the controlled microenvironments. Finally, we observed an upregulation of C3 in our glial cultures upon inflammation and further potentiation in cocultures, suggesting that C3 contributes to glial crosstalk in our culture model. Thus, the glial coculture model allows the successful mimicking of glial crosstalk and enhances our understanding of how microglia and astrocytes together contribute to neuroinflammation.

## Materials and Methods

### Differentiation of iPSC-derived microglia

The human iPSC line UTA.04511.WTs^27^ was used in this study for microglial differentiation. The line was derived and characterized at the Faculty of Medicine and Health Technology (MET), Tampere University, Finland with the approval of the Ethics Committee of Wellbeing Services County of Pirkanmaa (R08070). The hPSCs were acquired from a voluntary subject who had given written and informed consent. The project has received a supportive statement from the Wellbeing Services County of Pirkanmaa for the use of the named hPSC line in neuronal research (R20159). The iPSC line was expanded in feeder-free culture on 0.6 µg/cm^2^ recombinant human laminin-521 (LN521, BioLamina) in Essential8™ Flex media (E8 flex, Thermo Fisher Scientific), and passaged with TrypLE™ Select Enzyme and Defined Trypsin Inhibitor (DTI) (both from Thermo Fisher Scientific) in the presence of 10 µM Rock inhibitor (ROCKi, Y-27632, StemCell Technologies) as described previously^28^. The pluripotency of the iPSC line was monitored on a regular basis, and all cultures maintained normal karyotypes and were mycoplasma free.

The in-house-produced microglia were differentiated according to a previous publication^11^ with minor modifications (Fig. 1A). Briefly, on microglial differentiation Day 0, iPSCs were plated on Matrigel (Corning)-coated dishes at 9 000-20 0000 cells/cm^2^ and cultured under hypoxic conditions (5% O2, 5% CO2, 37 °C) until Day 4. On Days 0 and 1, the cells were cultured in E8 flex media supplemented with 5 ng/ml BMP4, 25 ng/ml activin A (both from Peprotech), 1 µM CHIR 99021 (Axon) and ROCKi (Day 0: 10 µM and Day 1: 1 µM). During Days 2 to 8, the cells were cultured in base media containing DMEM/F-12 without glutamine, 1X GlutaMAX, 543 mg/L sodium bicarbonate (all from Thermo Fisher Scientific), 14 µg/L sodium selenite, 64 mg/L L-ascorbic acid (both from Sigma–Aldrich) and 0.5% penicillin/streptomycin (P/S). On Days 2 and 3, the base media was supplemented with 100 ng/ml FGF2, 50 ng/ml VEGF (both from Peprotech), 10 µM SB431542 and 5 µg/ml insulin (both from Sigma–Aldrich). On Day 4, the cells were transferred to a normoxic incubator (5% CO2, 37 °C). From Day 4 until Day 8, the cells were cultured in base media supplemented with 50 ng/ml FGF2, 50 ng/ml VEGF, 50 ng/ml TPO, 50 ng/ml IL-6, 10 ng/ml SCF, 10 ng/ml IL-3 (all from Peprotech) and 5 µg/ml insulin (Sigma–Aldrich), and media changes were conducted daily. On Day 8, the floating erythromyeloid progenitor cells (EMPs) were collected and seeded at 64 000 cells/cm^2^ in ultra-low attachment dishes (Corning) in base media containing Iscove′s Modified Dulbecco′s medium (IMDM, Thermo Fisher Scientific), 10% heat-inactivated fetal bovine serum (FBS, Sigma–Aldrich) and 0.5% P/S supplemented with 5 ng/ml MCSF, 100 ng/ml IL-34 (both from Peprotech) and 5 µg/ml insulin. From Day 10 onward, the cells were cultured in primitive macrophage (PM) media containing IMDM supplemented with 10% FBS (Sigma–Aldrich), 0.5% P/S, 10 ng/ml MCSF and 10 ng/ml IL-34 (both from Peprotech). The media was changed every other day until final plating for the experiments was performed on Day 16. From Day 16 onward, half of the media was changed daily. For characterization, microglia were seeded at 45 00 cells/cm^2^ in 96-well plates (PerkinElmer) and cultured for 5 days prior to being fixed for immunocytochemistry. For RNA sample collection, the cells were plated at 41 700 cells/cm^2^ in 6-well plates (Thermo Fisher Scientific) and cultured for 6 days.

**Figure 1.**
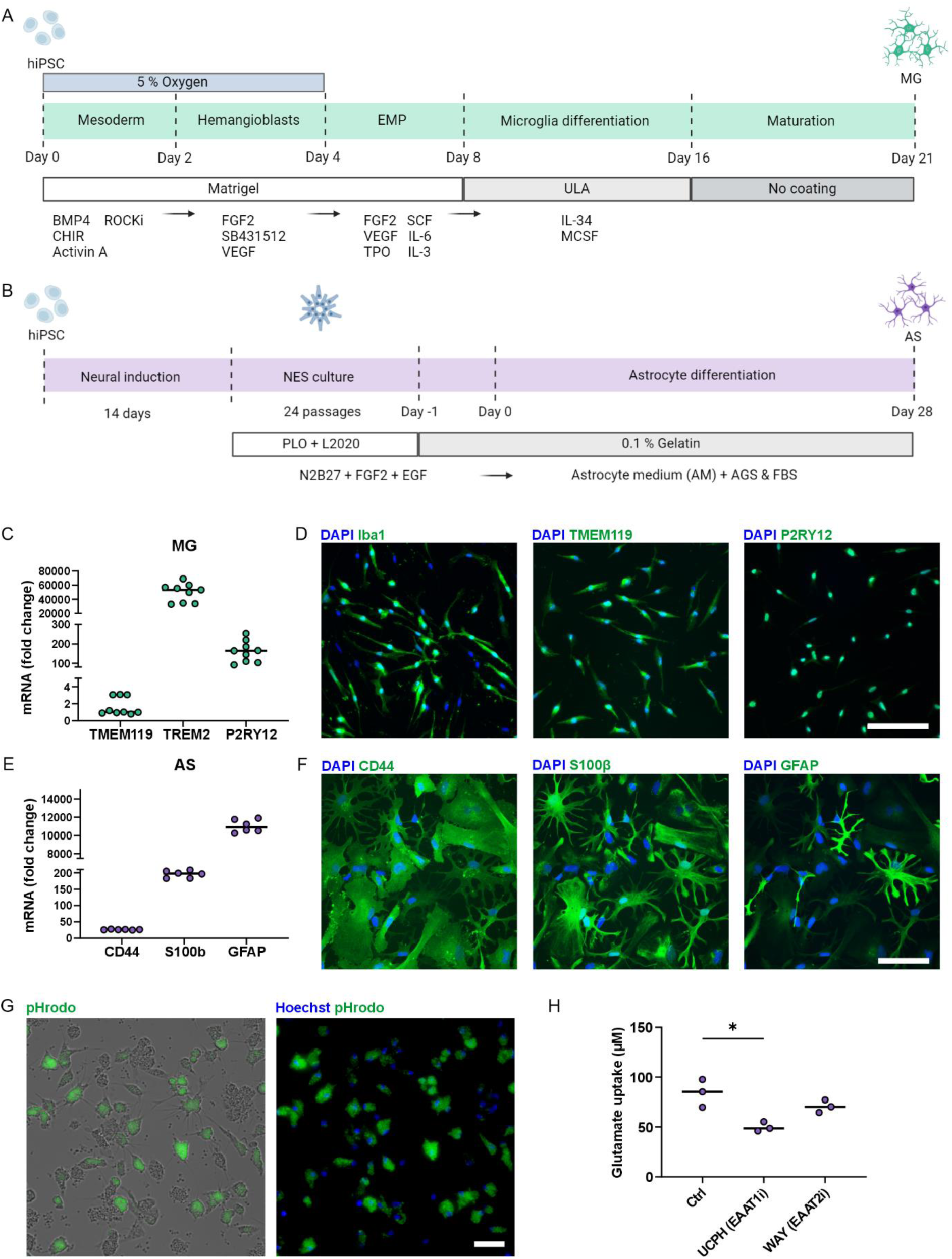
Characterization of differentiated hiPSC-derived microglia (MG) and astrocytes (AS). **A-B** Schematic timelines of (**A**) microglia differentiation and (**B**) astrocyte differentiation. Created in BioRender. Lotila, J. (2025) https://BioRender.com/j12k016. **C** RT– qPCR analysis of microglial expression of the genes *TMEM119*, *TREM2*, and *P2RY12*. n = 9 with 3 technical replicates, with 2 independent differentiations. **D** Representative immunofluorescence images of microglia characterized by the markers Iba1, TMEM119 and P2RY12. Scale bar, 100 µm. **E** RT–qPCR analysis of astrocyte expression of the genes *CD44*, *S100β* and *GFAP*. n = 6 with 3 technical replicates. **F** Representative immunofluorescence images of astrocytes characterized by the markers CD44, S100β and GFAP. Scale bar, 100 µm. **G** Representative live and fluorescence images of microglial phagocytosis of fluorescent pHrodo zymosan bioparticles. Scale bar, 50 µm. **H** Astrocyte glutamate uptake under control conditions and after inhibitor treatment (UCPH & WAY). n = 3 with 2 technical replicates. The data are presented as medians with individual values. *p< 0.05; Student’s t test.

### Differentiation of iPSC-derived astrocytes

Astrocyte differentiation in this study was conducted utilizing neuroepithelial stem cell (NES) line control 7 (NESC7)^29^, which was originally provided by the iPS Core Facility of Karolinska Institutet, Sweden. NESC7 cells were cultured and differentiated into astrocytes as described in a previous publication with minor modifications (Fig. 1B)^13^. Briefly, NESC7 cells were cultured in N2B27 media consisting of DMEM/F-12 GlutaMAX (Thermo Fisher Scientific), 1% N2 (Thermo Fisher Scientific), 0.1% B27 (Thermo Fisher Scientific), and 0.1% P/S supplemented with 10 ng/ml FGF2 (R&D Systems) and 10 ng/ml EGF (Sigma–Aldrich). The cells were cultured in culture flasks (T25 & T75, VWR) double-coated with 20 µg/ml poly-L-ornithine (PLO, Sigma–Aldrich) and 1:500 diluted murine Engelbreth-Holm-Swarm sarcoma derived laminin (L2020, Sigma–Aldrich). After passage 23–24, the cells were cultured in N2B27 media supplemented with a lower concentration of EGF (1 ng/ml). Full media change was conducted every other day, and on the days between media changes, growth factors were added to the culture media. The cells were passaged every 3–4 days at a ratio of 1:4–1:5 or at a density of 20 00–30 000 cells/cm^2^. Passaging was conducted using TrypLE™ Select Enzyme and DTI.

On Day −;1 of astrocyte differentiation, the NESs were passaged and seeded in 6-well plates coated with 0.1% gelatin (Thermo Fisher Scientific) at 30 000 cells/cm^2^ in N2B27 media supplemented with 10 ng/ml FGF2 and 1 ng/ml EGF. On Day 0, the cells were washed once with phosphate-buffered saline (PBS) before the addition of astrocyte media (AM) (ScienCell) supplemented with 2% FBS and 1% AGS (both from ScienCell) and 0.1% P/S. Media change with AM media was conducted every other day throughout the differentiation period. The cells were cultured without passaging until Day 6, after which the following passages were conducted within 1–2 days upon reaching 95% confluency. Passaging was conducted using TrypLE™ Select Enzyme and DTI. The cells were seeded at 30 000 cells/cm^2^ for all passages, and after the second passage, the cells were cultured in cell culture flasks (T25 and T75). On Day 28, differentiated astrocytes were plated for experiments. For characterization, astrocytes were plated at 30 000 cells/cm^2^ in 96-well plates (for immunocytochemistry) or 24-well plates (for RNA sample collection) (both from Thermo Fisher Scientific). The cells were cultured for 3 days before being fixed for immunocytochemical staining and for 4 days before RNA samples were collected.

### Establishment of microglia and astrocyte monocultures and cocultures

Microglia (MG) monocultures were prepared by seeding the cells in 96-well plates (PerkinElmer) at a density of 45 000 cells/cm^2^ on microglia differentiation Day 16 (Fig. 1A). The cells were allowed to mature for 5 days before the experiments were started.

Astrocyte (AS) monocultures were prepared by seeding the cells in PLO and LN521 double-coated 96-well plates at a density of 25 000 cells/cm^2^. Briefly, the well plates were coated with 0.1 mg/ml PLO for 1 hour at 37 °C, after which they were washed three times with sterile water and air-dried at room temperature (RT). The well plates were subsequentially coated with 15 µg/ml LN521 overnight at 4 °C. Astrocytes were plated in AM media. The media was changed to AM media supplemented with microglia growth factors 10 ng/ml MCSF and IL-34 (AM++) on the day after plating. Astrocytes were cultured for 7 days before the experiments were started. 50% media changes with AM++ were conducted daily.

Microglia and astrocyte (MG/AS) cocultures were prepared in PLO and LN521 double-coated 96-well plates (coated as described above). First, astrocytes were seeded in well plates at a density of 25 000 cells/cm^2^ in AM media on Day −;7. A full media change and switch to AM++ was conducted on the day after plating. On Day −;5, microglia were plated on top of the astrocytes at a density of 45 000 cells/cm^2^ on microglia differentiation Day 16. The cells were cultured together in AM++ media for 5 days before the experiments were started on Day 0. 50% of the media was changed daily.

### Design and fabrication of the microfluidic chip

An in-house-engineered microfluidic chip was used to culture microglia and astrocytes in their designated, interconnected compartments^19,30^. The utilized chip consists of two polydimethylsiloxane (PDMS) parts: a cell culture part and medium reservoir part (Supplementary Fig. 6). The cell culture part has three separate cell compartments that are interconnected by microtunnels. There are two microglia (MG) compartments on the sides (length = 3 mm, width = 4 mm) that are connected to the microglia/astrocyte (MG/AS) compartment in the middle (length = 5 mm, width = 4 mm) with 40 microtunnels (length = 250 µm, width = 10 µm, height = 3.5 µm). The microtunnels allow microglial migration between the compartments. The media reservoir part consists of three separate media chambers, enabling fluidic isolation between the cell compartments and the utilization of cell type-specific culture media.

The chips were fabricated from PDMS (SYLGARD 184, Dow Corning), and the cell compartments and microtunnels were treated with polyvinylpyrrolidone (PVP) (Sigma– Aldrich)^30^. The fabrication was conducted as described in a previous publication^31^ with few exceptions. The fabricated molds were treated with trichloro(1H,1H,2H,2H-perfluorooctyl)silane (Sigma–Aldrich) to improve the demolding process. The 4 mm thick PDMS sheets for the medium chambers were covered with a thin layer of dish soap before laser cutting to wash away the cut debris from the surfaces.

### Assembly of the microfluidic chip and cell culture

The assembly of the microfluidic chips was conducted as described previously^19^. Briefly, HCl-cleaned ∅24 mm glass coverslips were coated with 0.25 mg/ml PLO at 37 °C for 1.5 h. Subsequently, the coverslips were washed three times with sterile H2O, allowed to air-dry at RT, and stored at 4 °C. PVP-coated microfluidic chips were sterilized by immersion in 70% ethanol and air-dried at RT prior to assembly. Thereafter, the microfluidic chips were manually attached to PLO-coated glass coverslips and the cell compartments were coated with 30 µg/ml LN521 overnight at 4 °C.

Microglia were seeded in their designated side compartments at a density of 25 000 cells/cm^2^ on microglia differentiation Day 16 (Day −;7). The microglia were cultured on the chip for 5 days to allow maturation prior to seeding astrocytes. During this period, PM media was used for the MG compartments, and AM media was used for the middle MG/AS compartment. On Day −;2, astrocytes were seeded into the MG/AS compartment at a density 45 000 cells/cm^2^ in AM++ media supplemented with microglial growth factors. The cells were cultured together in the microfluidic chip for two days before the experiments were started on Day 0. Fresh media for the media chambers was changed daily throughout the culture without disturbing the cell compartments.

### Fluidic isolation on the microfluidic chip

To demonstrate fluidic isolation between the cell compartments on the chip, FITC-conjugated dextran particles (15–25 kDa) (TdB Consultancy) were used. The particles (50 µM) were added to the middle MG/AS compartment, and their diffusion was evaluated for 24 h at 37 °C. The diffusion of the dextran particles into the outer MG compartments was visualized with an Olympus IX51 microscope equipped with an Olympus DP30BW camera (Olympus Corporation). To quantify the amounts of dextran particles in the compartments after 24 h, the absorbance of the medium samples at 490 nm was measured using a NanoDrop 1000 spectrophotometer (Thermo Fisher Scientific).

### Stimulation of inflammatory activation

Inflammatory activation of MG and AS monocultures and MG/AS cocultures was stimulated for 24 hours using three different stimulants: 100 ng/ml lipopolysaccharide (LPS) (Sigma– Aldrich), a combination of 10 ng/ml TNF-α/IL-1β (TI) (both from Peprotech), or 20 ng/ml interferon (IFN)-γ (Peprotech) in respective culture medias (PM or AM++). After 24 h of treatment, viability staining was performed, the media were collected for secretion analysis, and the cultures were fixed for immunocytochemical staining.

The stimulation of inflammatory activation in the microfluidic chip cocultures was conducted in a similar manner. On Day 0, culture media (PM or AM++) containing 100 ng/ml LPS, a combination of 10 ng/ml IL-1β/TNF-α (TI), or 20 ng/ml IFN-γ was added to each cell compartment and media chamber. For the migration studies, the cells were additionally stimulated with 100 µM adenosine diphosphate (ADP) (Sigma–Aldrich). The cells were treated for 24 hours, after which culture media was collected from media chambers of both the MG and MG/AS compartments. Media from MG compartments of the same chip were pooled, but each microfluidic chip was collected separately. The cells were fixed for immunocytochemical staining.

### Immunocytochemistry

Immunocytochemistry (ICC) of conventional cultures was performed by first fixing the cultures for 15 minutes with 4% paraformaldehyde in PBS. The fixed samples were then blocked for 45 minutes with 10% normal donkey serum (NDS), 0.1% Triton X-100, and 1% bovine serum albumin (BSA) in PBS at RT. Primary antibody solutions were diluted in 1% NDS, 0.1% Triton X-100, and 1% BSA in PBS and incubated overnight at 4 °C. The secondary antibodies were diluted in 1% BSA in PBS and incubated for 1 h at RT. The samples were then mounted with ProLong™ Gold Antifade Mountant with DAPI (Thermo Fisher Scientific). ICC on microfluidic chips was performed as described in a previous publication^19^. As a minor modification, the microfluidic chips were washed three times with 4’,6-diamidino-2-phenylindole (DAPI) diluted 1:5 000 in PBS after secondary antibody incubation. The primary and secondary antibodies used are listed in Supplementary Table 1. Images were acquired with an Olympus IX51 inverted fluorescence microscope equipped with a Hamamatsu ORCA-Flash4.0 LT + sCMOS camera (Olympus Corporation), a DMi8 inverted microscope (Leica), and an LSM780 laser-scanning confocal microscope equipped with a Quasar spectral GaAsP detector (all from Carl Zeiss). Image analysis was performed with CellProfiler and CellProfiler Analyst software^32^.

### RNA isolation and quantitative PCR

RNA was extracted from astrocytes, microglia, NESs and iPSCs utilizing a NucleoSpin RNA kit (Macherey-Nagel). The purity and concentration of RNA were measured with a NanoDrop 1000 (Thermo Fisher Scientific). The RNA was converted to cDNA with a High Capacity cDNA Reverse Transcription Kit (Thermo Fisher Scientific). The gene expression levels of the astrocytic markers *CD44* (Hs01075864_m1), *S100β* (Hs00902901_m1) and *GFAP* (Hs00909236_m1) and the microglial markers *TMEM119* (Hs01938722_u1), *P2RY12* (Hs00224470_m1) and *TREM2* (Hs00219132_m1) were analyzed with TaqMan assays via the ABI QuantStudio 12K Flex Real-Time PCR System (Thermo Fisher Scientific). The samples were run in triplicate. The data were analyzed via the ΔΔCt method. Astrocyte data were normalized to those of the housekeeping gene *GUSB* (Hs00939627_m1) and NES samples. The microglia results were normalized to housekeeping gene *GAPDH* (Hs99999905_ m1) and the iPSC sample.

### Glutamate uptake assay

Glutamate uptake by differentiated astrocytes was analyzed with a glutamate assay kit (Abcam). Astrocytes were plated at 100 000 cells/cm^2^ in a 24-well plate (Thermo Fisher Scientific) and cultured for 4 days in AM medium before the assay was started. First, the cells were washed twice with PBS (with Ca^++^ and Mg^++^), followed by a 30 min incubation at 37 °C with 1:10 diluted DMSO (control, Sigma–Aldrich), 1.5 mM UCPH-101 (EAAT-1 inhibitor, Abcam), or 1 mM WAY213613 (EAAT-2 inhibitor, Tocris) in Hank’s balanced salt solution (with Ca^++^ and Mg^++^) (HBSS, Gibco). After incubation, HBSS containing 100 µM glutamate was added to the cells, which were subsequently incubated for 60 min at 37 °C. The cells were then washed three times with PBS and lysed with assay buffer. The concentration of glutamate in the cell lysates was analyzed following the manufacturer’s protocol.

### Mesoscale and ELISA

The secretion of cytokines and chemokines (TNF-α, IL-1β, IL-6, GM-CSF, IL-10, CCL2, CXCL5, CXCL8, and CXCL10) from the collected culture medium was measured using a U-PLEX Custom Biomarker Group 1 (human) Assay (Meso Scale Diagnostics) according to the manufacturer’s protocol. Secretion of C3 was measured using a R-PLEX Human Complement C3 Assay (Meso Scale Diagnostics). The plates were run using a MESO QuickPlex SQ 120 instrument and the results were analyzed with DISCOVERY WORKBENCH® software (v4.0) (Meso Scale Diagnostics). If the measured values were below the detection limit, the values were set to 0. Values above the maximum detection limit were set to the maximum detectable value. The levels of IL-6 in the medium were measured with a human IL-6 uncoated ELISA kit (Thermo Fisher Scientific) according to the manufacturer’s protocol. The absorbance was measured at 450 nm (Wallac Victor 1420).

### Phagocytosis assay

The phagocytic capacity of differentiated microglia was investigated using pHrodo™ Green Zymosan Bioparticles™ Conjugate for Phagocytosis (P35365, Thermo Fisher Scientific). pHrodo bioparticles were added to cells at 100 µg/ml in Opti-MEM (Thermo Fisher Scientific) supplemented with the microglial growth factors IL-34 and MCSF (both 10 ng/ml). After 6 h of incubation, the nuclei were stained with Hoechst 33342 (1:1 000, Thermo Fisher Scientific), and the cells were imaged immediately with an Olympus IX51 fluorescence microscope equipped with a Hamamatsu ORCA-Flash4.0 LT + sCMOS camera.

### Viability staining

The viability of the cultured cells after inflammatory stimulation was analyzed via a viability/cytotoxicity kit for mammalian cells (Thermo Fisher Scientific). The cultures were incubated for 30 min at 37 °C with green fluorescent calcein-AM (0.5 µM) for the detection of live cells and red fluorescent ethidium homodimer-1 (EthD-1) (0.5 µM) for the detection of dead cells. The samples were imaged with an Olympus IX51 microscope equipped with a Hamamatsu ORCA-Flash4.0 LT + sCMOS camera.

### Statistical analysis

The normality of the data was determined with the Shapiro–Wilk test. Statistical analysis of normally distributed data was performed with Student’s t test or one-way ANOVA with Tukey’s post hoc comparison. The nonparametric Mann–Whitney U test was used for nonnormally distributed data. A p value < 0.05 was considered statistically significant. Statistical significance is denoted as * p < 0.05, ** p < 0.01, and *** p < 0.001. All the statistical tests were performed with IMB SPSS Statistics software (version 29.0.1.0).

## Results

### Generation of cocultures of human iPSC-derived microglia and astrocytes exhibiting cell type-specific markers and functions

We used well-established published protocols to differentiate iPSC-derived microglia^11^ and astrocytes^13^ (Fig. 1A and B). Successful microglial differentiation was confirmed by the gene expression of the typical markers *TMEM119*, *TREM2*, and *P2RY12* (Fig. 1C) and positive immunocytochemical staining for Iba1, TMEM119, and P2RY12 (Fig. 1D). Differentiated astrocytes, in turn, expressed the astrocyte-specific markers CD44 and S100 calcium-binding protein beta (S100β) and the mature astrocyte marker glial fibrillary acidic protein (GFAP) at both the gene (Fig. 1E) and protein levels (Fig. 1F). Both cell types exhibited key functions: microglia, through the phagocytosis of pHrodo zymosan bioparticles (Fig. 1G), and astrocytes, via glutamate uptake, which was primarily mediated by the glutamate transporter EAAT1, as shown by reduced uptake after treatment with the inhibitor UCPH-101 (Fig. 1H). Therefore, both cell types displayed cell type-specific characteristics and performed essential functions.

To study the immune responses of both cell types, we prepared microglia and astrocyte monocultures (MG and AS) and microglia/astrocyte cocultures (MG/AS) (Fig. 2A and B). Equal amounts of microglia and astrocytes were plated for both culture setups to enable comparison. Quantification revealed that after six days, the stabilized cocultures consisted of 11% microglia and 89% astrocytes (Fig. 2E). Viability staining with calcein-AM and EthD-1 demonstrated high viability of the established cultures, which was also indicated by negative immunocytochemical staining with the apoptotic marker cleaved caspase-3 (Supplementary Fig. 1).

**Figure 2.**
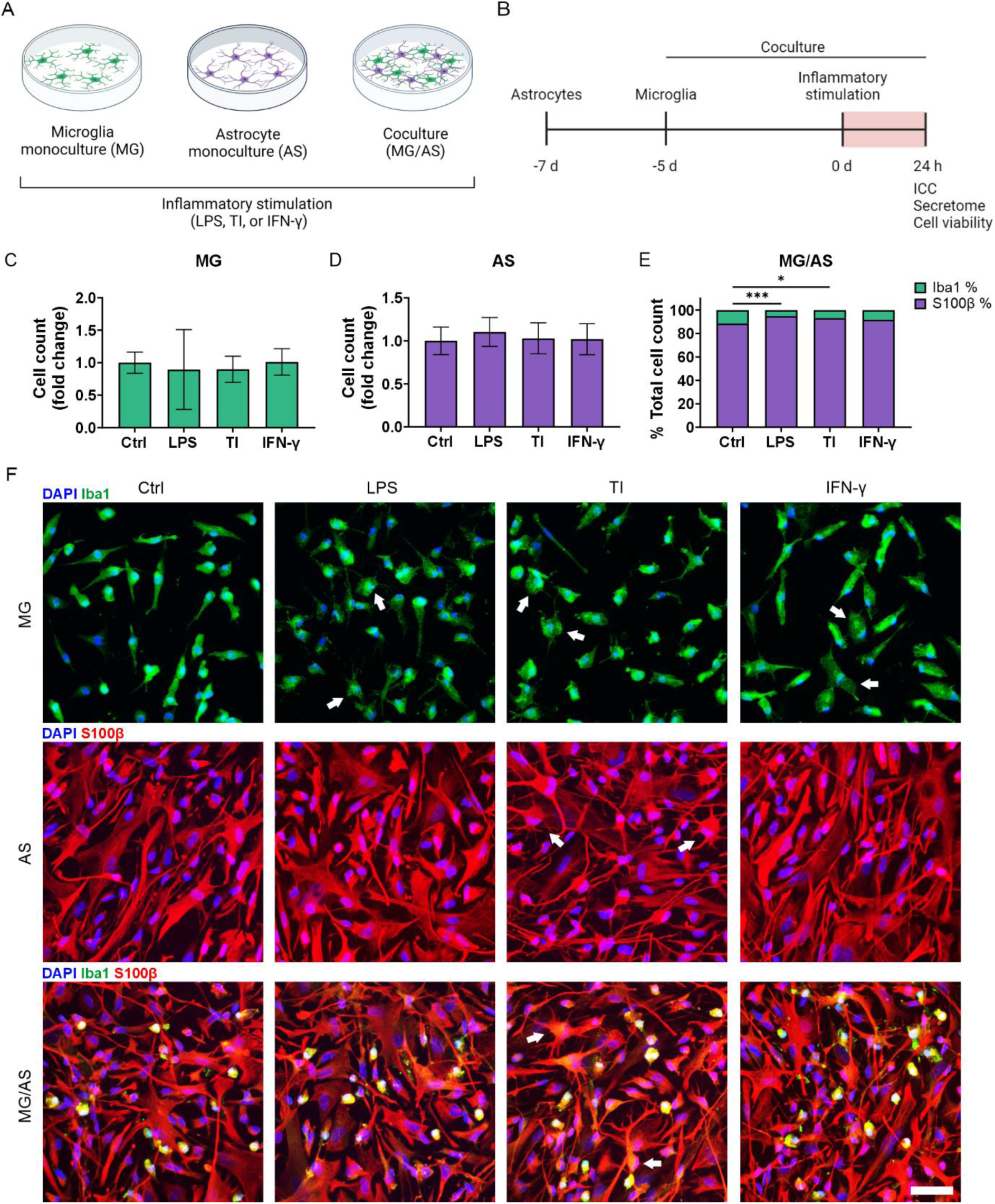
Inflammatory stimulation of established glial monocultures (MG, AS) and cocultures (MG/AS). **A-B** Schematic figures of (**A**) the experimental setup and (**B**) timeline. Created in BioRender. Lotila, J. (2025) https://BioRender.com/j12k016. **C-D** Quantified cell amounts of monocultures are presented as the fold change. n = 18 images per condition; images from 2 experiments. The data are presented as the means ± SDs. **E** Quantification of Iba1- and S100β-positive cells in cocultures presented as a percentage of the total cell count. n = 18 images per condition; images from 2 experiments. The data are presented as the means. **F** Representative immunofluorescence images showing observable morphological changes following inflammatory activation in both monocultures and cocultures. The white arrows indicate the observed morphological changes. Scale bar, 50 µm. *p< 0.05, **p< 0.01, ***p< 0.001; one-way ANOVA with Tukey’s post hoc comparison.

### Neuroinflammatory stimulation induces characteristic morphological changes without affecting cell viability

To study the immune responses of microglia and astrocytes, we stimulated established cultures for 24 hours with the following inflammatory stimulants: LPS, TNF-α/IL-1β, or IFN-γ (Fig. 2A and B). First, we investigated whether stimulation had an effect on the overall number of cells and their proliferation. MG and AS monocultures presented no significant alterations in cell numbers following stimulation (Fig. 2C and D). The proportion of microglia in cocultures was more variable after stimulation and decreased with both LPS (5% vs. 11%, p<0.001) and TNF-α/IL-1β stimulation (7% vs. 11%, p=0.019) compared with the control conditions (Fig. 2E). Immunocytochemical staining of the proliferation marker Ki-67 revealed reduced proliferation in all the stimulation groups in the MG monocultures compared with the control (p<0.001), which could partially explain the small reduction in cell numbers (Supplementary Fig. 2).

We further investigated the potential effects of inflammatory stimulation on the morphology and viability of the prepared cultures. Observation of cell morphology revealed that all stimulants increased hypertrophy in MG monocultures, indicating transition to an activated state (Fig. 2F). TNF-α/IL-1β stimulation induced increased ramification in AS monocultures, but LPS or IFN-γ stimulation did not have observable effects. While astrocytes in cocultures demonstrated comparable morphological changes in response to TNF-α/IL-1β stimulation, microglial morphology was already more amoeboid under control conditions, and consequently did not demonstrate similar observable morphological changes as those in monocultures. Finally, viability staining with calcein-AM and EthD-1 confirmed high viability in all stimulated cultures (Supplementary Fig. 3), which was also supported by negative immunocytochemical staining for the apoptotic marker cleaved caspase-3 (Supplementary Fig. 4). In summary, inflammatory stimulation of glial cultures induced typical observable morphological changes in both microglia and astrocytes, indicating inflammatory activation with no detrimental effects on cell viability.

### Stimulation with LPS and TNF-α/IL-1β induces cell type-specific responses in microglia and astrocytes and triggers glial crosstalk in cocultures

We analyzed the inflammatory secretome with a multiplex assay to investigate the effects of different stimulants as well as glial crosstalk on inflammatory responses in the studied cultures (Fig. 3A). LPS stimulation induced a strong inflammatory response in MG monocultures, resulting in increased secretion of chemokines (CXCL5, CCL2 and CXCL8) and cytokines (IL-6, IL-10, TNF-α, and IL-1β) compared to control level (Fig. 3B). Similarly, TNF-α/IL-1β stimulation induced a strong inflammatory response in AS monocultures, resulting in prominent secretion of chemokines (CXCL10, CXCL5, CCL2, and CXCL8) and cytokines (GM-CSF and IL-6) as well as low secretion of IL-10 (Fig. 3B). Thus, microglia and astrocytes demonstrated cell type-specific responses to LPS and TNF-α/IL-1β stimulation, respectively, which was further validated by statistical tests conducted between the groups (Fig. 3C and D). IFN-γ stimulation, in turn, generated more moderate responses in both cell types, inducing the secretion of CXCL10, CCL2, CXCL8, and IL-1β in MG monocultures and low secretion of CCL2 and IL-6 in AS monocultures (Fig. 3B, Supplementary Fig. 5A).

**Figure 3.**
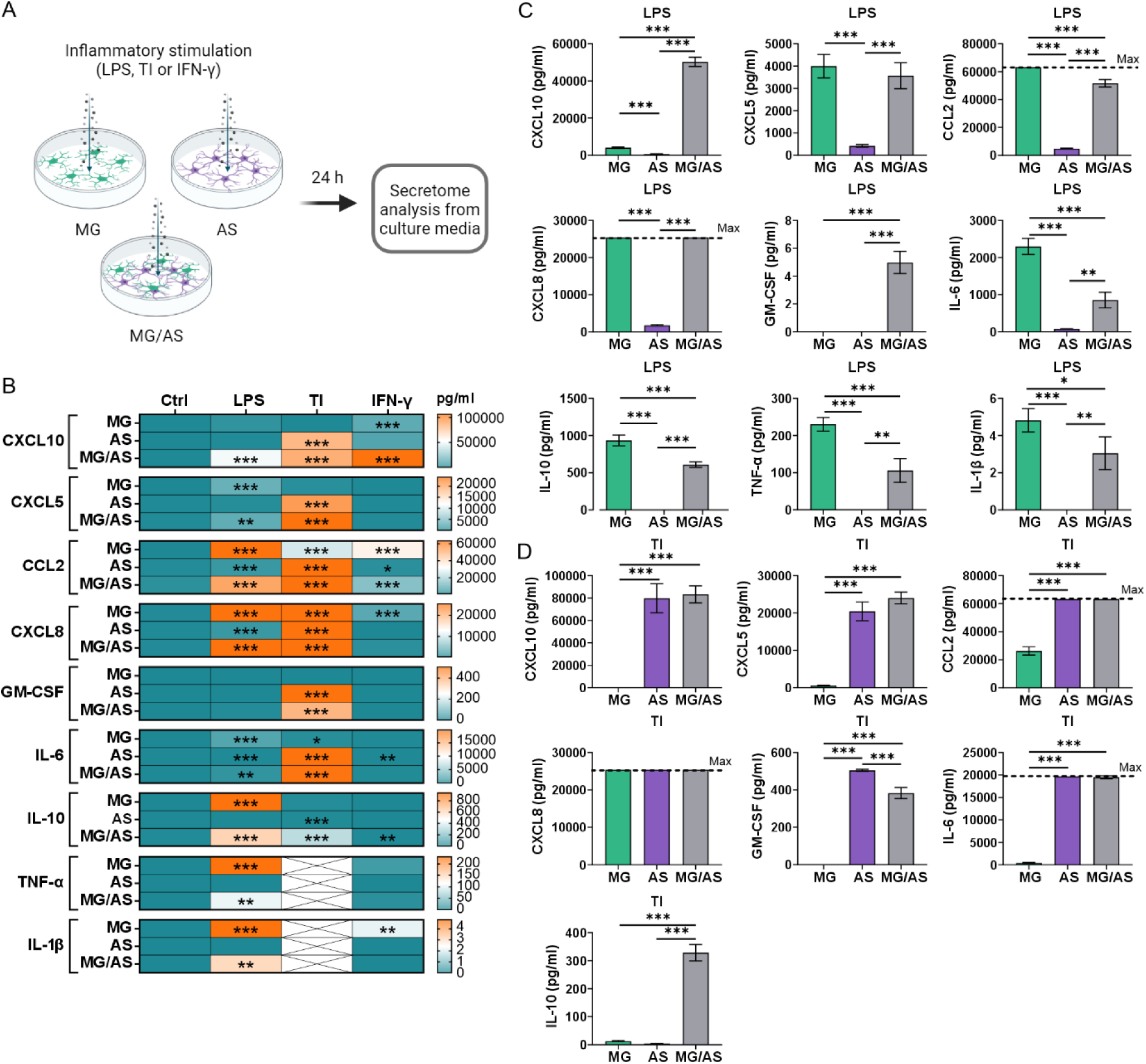
Secretion of inflammatory factors in conventional cultures following inflammatory stimulation. **A** Schematic figure of inflammatory stimulation. Created in BioRender. Lotila, J. (2025) https://BioRender.com/j12k016. **B** Stimulated cultures secreted a wide range of inflammatory factors as a result of inflammatory activation. Asterisks indicate statistical significance in comparison with the control. n = 2–3, with 2 technical replicates; the data are representative of 2 experiments. The data are presented as the means. **C** Comparison of secretion levels between different cultures stimulated with LPS or **D** TNF-α/IL-1β (TI). n = 3, with 2 technical replicates; the data are representative of 2 experiments. The data are presented as the means ± SDs. *p< 0.05, **p< 0.01, ***p< 0.001; one-way ANOVA with Tukey’s post hoc comparison. The dotted line indicates the maximum detection of the analyte.

Next, we wanted to understand how the inflammatory responses of microglia and astrocytes are modulated in cocultures. While MG/AS cocultures exhibited patterns of cell type-specific inflammatory activation similar to those observed in monocultures (Fig. 3B), the secreted levels of several chemokines and cytokines were significantly altered, indicating reciprocal modulation of inflammatory responses between the cell types. Compared with MG monocultures, LPS stimulation of MG/AS cocultures induced greater secretion of CXCL10 and GM-CSF and lower secretion of CCL2, IL-6, IL-10, TNF-α, and IL-1β (Fig. 3C). Interestingly, elevated levels of CXCL10 and GM-CSF were detected in TNF-α/IL-1β-stimulated AS monocultures but not in LPS-stimulated MG monocultures (Fig. 3B). These findings suggest that their increase in LPS-stimulated MG/AS cocultures indicates inflammatory activation of astrocytes through crosstalk with LPS-activated microglia.

In contrast, TNF-α/IL-1β stimulation induced greater secretion of IL-10 and lower secretion of GM-CSF in MG/AS cocultures than in AS monocultures (Fig. 3D). Similarly, since IL-10 was secreted primarily by LPS-stimulated MG monocultures, its secretion in TNF-α/IL-1β-stimulated MG/AS cocultures suggests that inflammatory activation of microglia occurs through crosstalk with inflammatory astrocytes. Thus, while LPS and TNF-α/IL-1β induced strong cell type-specific inflammatory responses in monocultures, the secretion of various chemokines and cytokines was modulated in cocultures, demonstrating glial crosstalk.

### Microfluidic coculture platform facilitates the study of microglial migration and glial activation within distinct inflammatory microenvironments

To better model glial crosstalk and the effects of inflammatory microenvironments on glial functions, we established a microfluidic coculture chip platform (Fig. 4A, Supplementary Fig. 6). The chip platform consisted of three sequential cell compartments interconnected by microtunnels^19,30,31^. This enabled the culture of microglia and astrocytes in their separate dedicated compartments (Fig. 4B and C), and the microtunnels facilitated spontaneous microglial migration toward the astrocyte compartment (MG/AS C) (Fig. 4C and D), which increased over an extended culture period (Fig. 4E). Each compartment was provided with its own media chamber to allow the culture of microglia and astrocytes in cell type-specific media as well as the establishment of separate inflammatory microenvironments to study inflammatory crosstalk. Fluidic isolation between the different compartments was confirmed using FITC-dextran particles of a size comparable to that of secreted cytokines (20 kDa) (Fig. 4F and G). The results revealed efficient fluidic isolation with no diffusion of fluorescent FITC-dextran particles between the compartments over a 24-hour test period.

**Figure 4.**
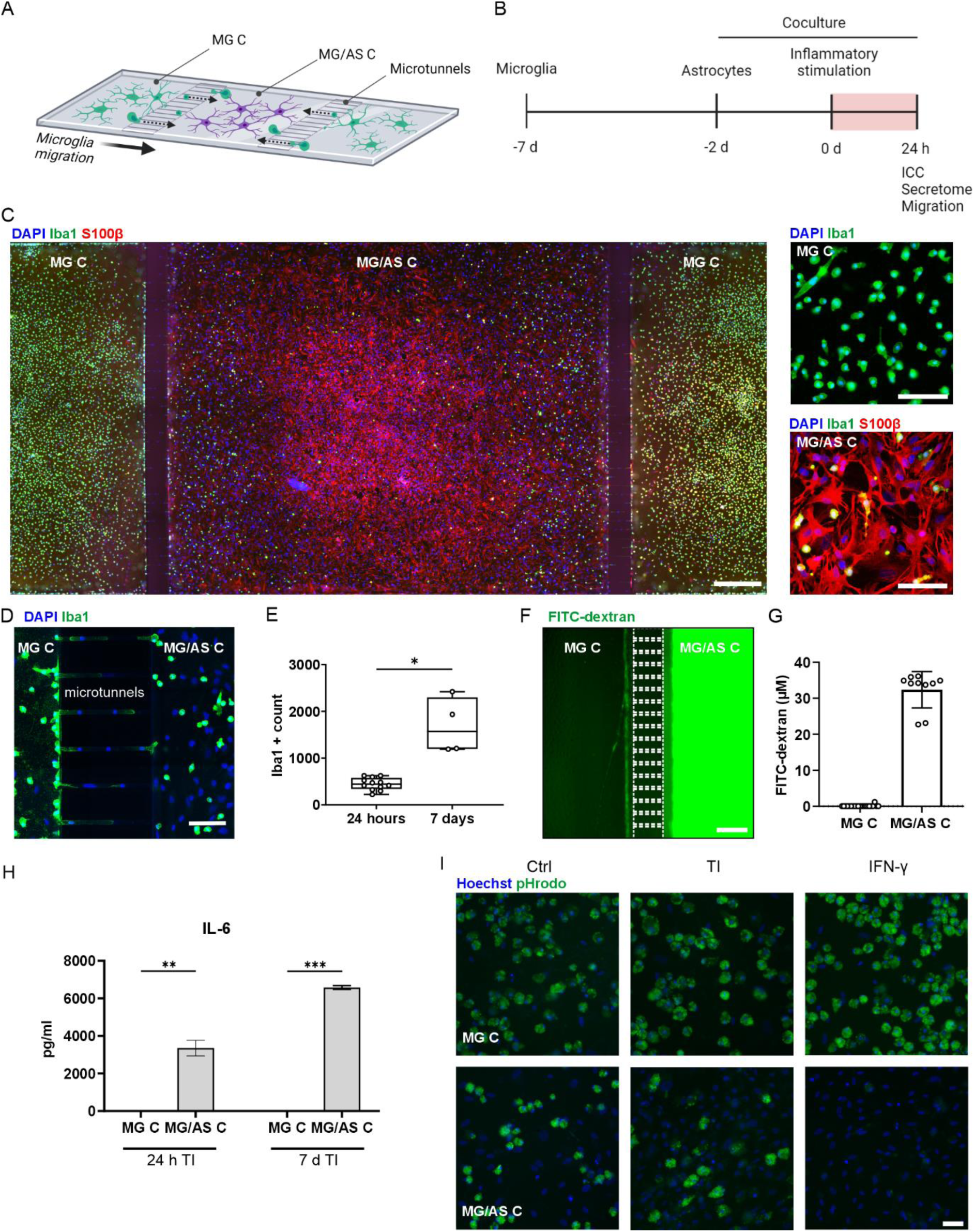
Microglia (MG) and astrocyte (AS) coculture in the microfluidic chip. **A** Schematic figure of the microfluidic chip design and coculture in the chip. **B** Timeline of the experimental setup. Created in BioRender. Lotila, J. (2025) https://BioRender.com/j12k016. **C** Representative immunofluorescence image of the cells cultured in the compartmentalized chip. Scale bar, 500 µm. Close-up images of microglia in MG C and microglia and astrocyte cocultures in MG/AS C. Scale bar, 100 µm. **D** Representative immunofluorescence image showing close-up of Iba1-positive microglia migrating through the microtunnels. Scale bar, 100 µm. **E** Microglia demonstrated spontaneous migration toward the MG/AS C, which increased over time. n(24 hours) = 12, n(7 days) = 4. The data are presented as boxplots showing independent values and whiskers indicating min and max. *p< 0.05; Student’s t test. **F** Representative fluorescence image of FITC-dextran particles in the chip after 24 hours. Scale bar, 250 µm. **G** Measurement of the concentration of FITC-dextran particles from different compartments after 24 hours. n = 10–21. The data are presented as the means ± SDs. **H** IL-6 secretion measured with ELISA in different compartments after 24 hours and 7 days of selective MG/AS C inflammatory stimulation with TNF-α/IL-1β (TI). n = 2 with 2 technical replicates. The data are presented as the means ± SDs. **p<0.01, ***p<0.001; Student’s t test. **I** Representative fluorescence images showing microglial phagocytosis of fluorescent pHrodo bioparticles in different compartments after 6 d of selective MG/AS C inflammatory stimulation with TNF-α/IL-1β (TI) or IFN-γ. Scale bar, 50 µm.

To further confirm the establishment of separate microenvironments within the chip, we stimulated only MG/AS C with TNF-α/IL-1β for 24 h and 7 d and measured the secretion of IL-6 from both MG C and MG/AS C. Elevated IL-6 was measured only within the stimulated compartment, ensuring that it did not diffuse into the neighboring MG compartment (Fig. 4H). In addition, the functionality of migrating microglia was evaluated via a pHrodo phagocytosis assay. We confirmed that microglia migrating to the MG/AS C performed phagocytic functions in the basal state (Fig. 4I). However, after targeted proinflammatory stimulation of MG/AS C with IFN-γ, we observed compromised phagocytic function in the inflammatory microenvironment, whereas microglia in unstimulated MG compartments remained unaffected. Thus, the microfluidic chip platform demonstrated successful coculturing of microglia and astrocytes with intercompartmental microglial migration and fluidically isolated compartments, allowing investigations of glial activation within distinct microenvironments.

### The chemoattractant ADP increases microglial migration in a microfluidic chip platform

To explore how inflammatory stimulation affects microglial migration in the chip, we investigated the number of Iba1-positive cells in the MG/AS C after 24 hours of stimulation of all compartments (Fig. 5A and B). In addition, we stimulated the compartments with ADP, a known stimulant of microglia migration^33^. First, we studied the effects of stimulants on microglia cultured in a microfluidic chip without astrocytes (MG). Microglia increased their migration toward MG/AS C after 24 hours of ADP stimulation (p= 0.01) (Fig. 5C). There was also a modest increase in migration in response to TNF-α/IL-1β, but this increase was not statistically significant. In contrast, microglial migration toward MG/AS C in the presence of astrocytes (MG/AS) was more variable than that of microglia cultured alone, and no statistically significant differences between the stimulation groups were detected, although a trend of increased migration following the addition of ADP was noted (Fig. 5D and E). Thus, while the presence of astrocytes and proinflammatory challenge did not result in major changes in microglial migration, we were able to observe an increase in microglial migration upon exposure to the ADP stimulus.

**Figure 5.**
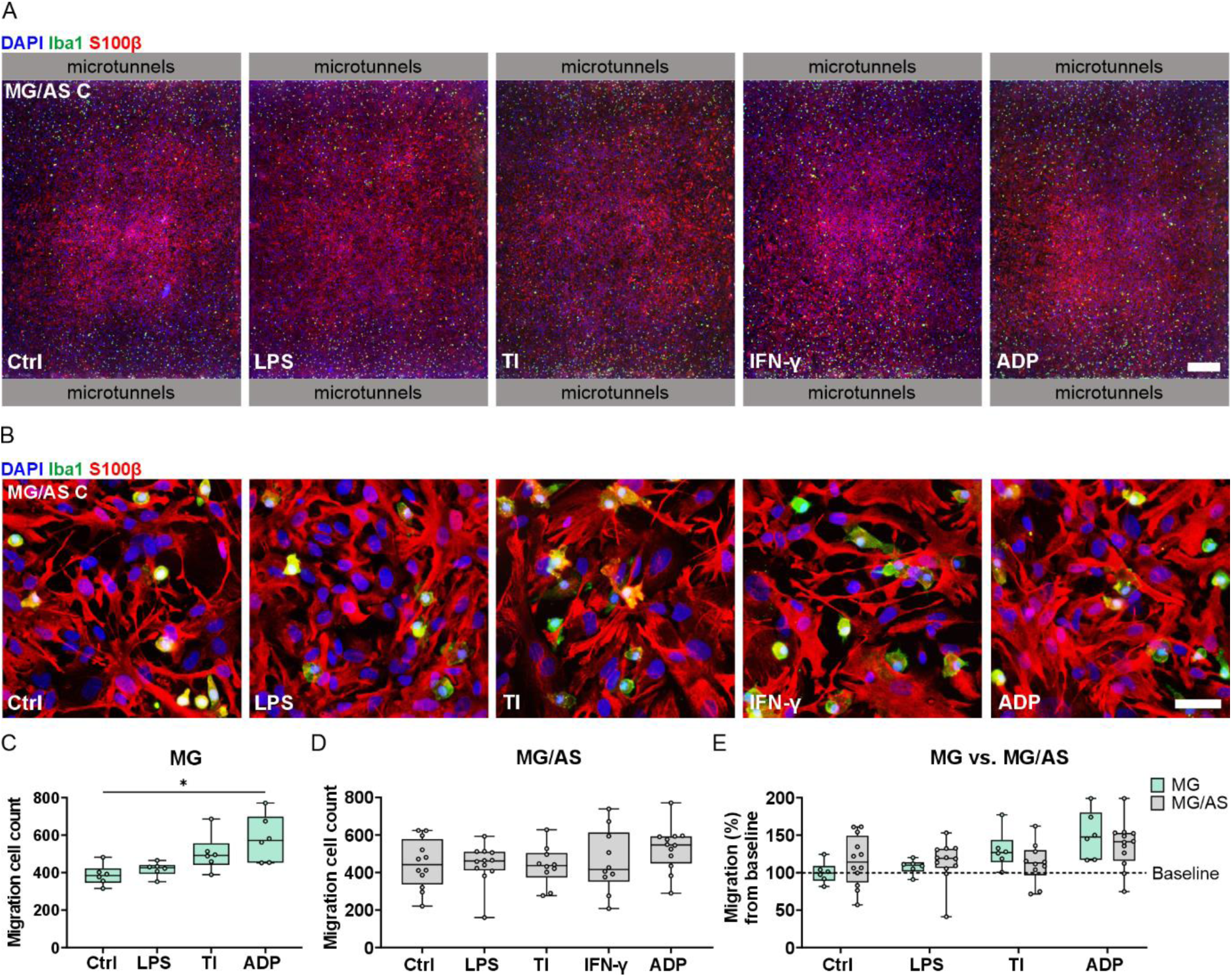
Microglial migration in microfluidic chip cocultures after 24 hours of inflammatory stimulation. **A** Representative immunofluorescence images of the MG/AS C in coculture chips after 24 hours of inflammatory stimulation. Scale bar, 500 µm. **B** Representative immunofluorescence images showing close-up of cocultures in the MG/AS C. Scale bar, 50 µm. **C** Cell count of microglia that migrated into the MG/AS C without astrocytes (MG) after 24 hours of stimulation. n = 6 with 3 chips in each group; migration through microtunnels analyzed separately for both sides. Microglial migration significantly increased following ADP stimulation. **D** Count of microglia that migrated into the MG/AS C in coculture chips (MG/AS) after 24 hours of stimulation. n = 10–12 with 5–6 chips in each group; migration through microtunnels analyzed separately for both sides. No significant alterations in migration were detected between the control and stimulated groups. **E** Migration of MG chips and MG/AS coculture chips after 24 hours of stimulation as a percentage of the MG baseline. n(MG) = 6 with 3 chips in each group, n(MG/AS) = 10–12 with 5–6 chips in each group. The data are presented as boxplots showing independent values and whiskers indicating min and max. *p< 0.05; Mann–Whitney U test.

### Inflammatory stimulation of microfluidic chip cocultures successfully recapitulates glial crosstalk with unique responses

To verify that we can reproduce the observed inflammatory activation and crosstalk of microglia and astrocytes in chip microenvironments, we stimulated all compartments and analyzed the secretion of inflammatory factors from the MG C and MG/AS C media chambers (Fig. 6A). To enable comparisons between conventional culture and chip results, we also included chip cultures that contained only astrocytes in their designated compartment (AS C). Compared with conventional cultures, the cultures on a microfluidic chip presented considerably lower levels of secretion, reflecting overall smaller cell quantities (Fig. 6B–D). However, as in conventional monocultures (Fig. 3B), both LPS and TNF-α/IL-1β stimulation induced strong cell type-specific immune responses in the corresponding compartments, MG C and AS C (Fig. 6B). Accordingly, IFN-γ stimulation induced minor responses in both cell types (Fig. 6B, Supplementary Fig. 5B).

**Figure 6.**
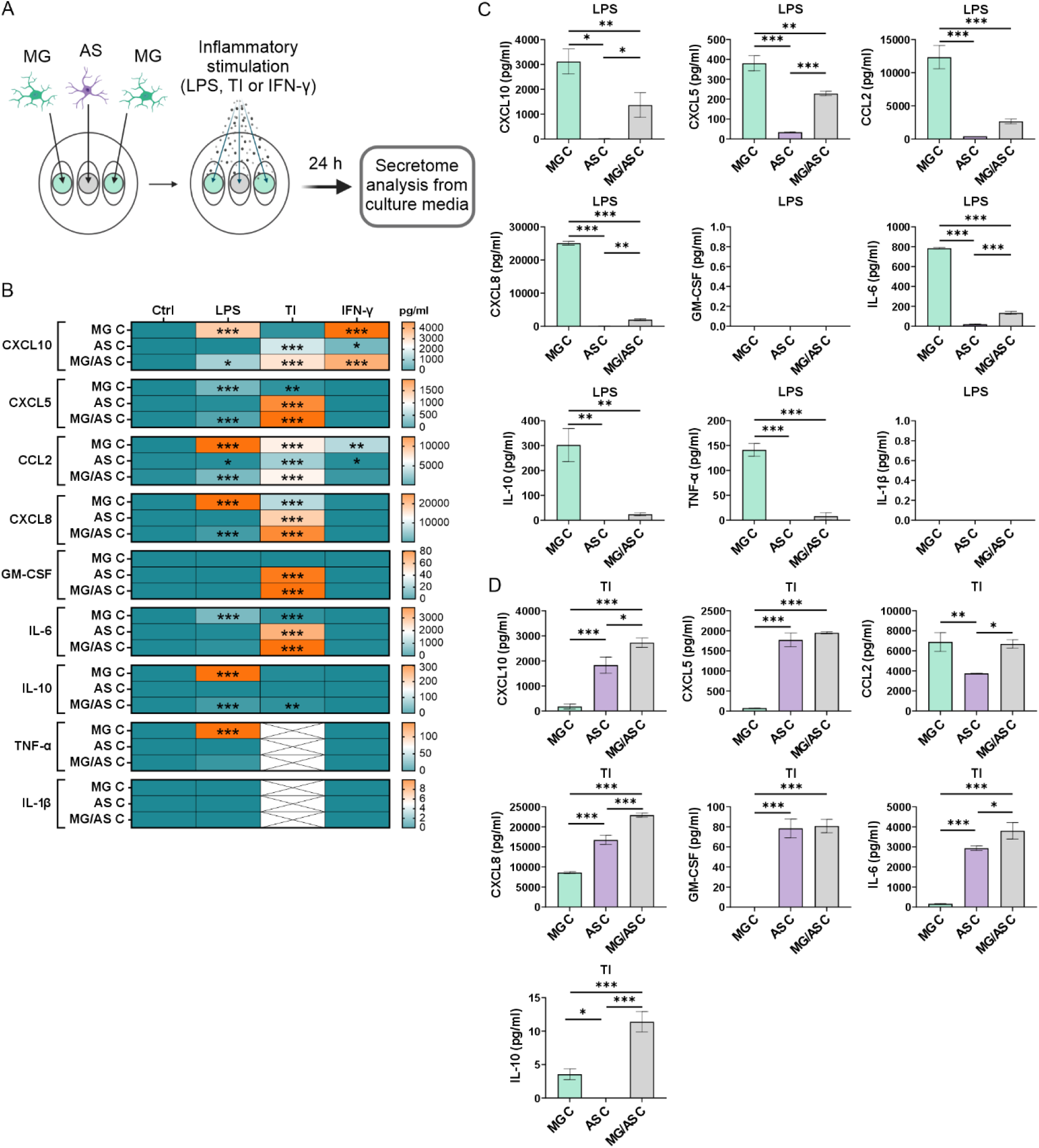
Secretion of inflammatory factors in microfluidic chip microglia and astrocyte cocultures. **A** Schematic figure of the coculture setup and inflammatory stimulation. Created in BioRender. Lotila, J. (2025) https://BioRender.com/j12k016. **B** Heatmap of the secretion of inflammatory cytokines and chemokines in the chip. Asterisks indicate statistical significance in comparison with the control. **C** Comparison of secretion levels between different cultures stimulated with LPS or **D** TNF-α/IL-1β (TI). The dotted line indicates the maximum detection of the analyte. n = 2–3, with 2 technical replicates for all data; the data are representative of 2 experiments except for AS C, which was repeated once. The data are presented as the means ± SDs. *p< 0.05, **p< 0.01, ***p< 0.001; one-way ANOVA with Tukey’s post hoc comparison.

Compared with conventional MG/AS cocultures, chip cocultures of MG/AS C produced similar patterns of LPS-induced responses, confirming the contribution of migrated microglia to inflammatory activation (Fig. 3B and C and Fig. 6B and C). Furthermore, TNF-α/IL-1β stimulation increased IL-10 secretion in MG/AS C, similar to what was observed in conventional MG/AS cocultures (Fig. 3D and Fig. 6D). Interestingly, compared with AS C monocultures, MG/AS C cocultures presented mostly enhanced responses to TNF-α/IL-1β stimulation, resulting in increased secretion of CXCL10, CCL2, CXCL8, IL-6, and IL-10 (Fig. 6D), which was not observed in conventional MG/AS cocultures (Fig. 3D). Overall, we have demonstrated that microglia and astrocytes cocultured in the microfluidic chip platform reproduce the inflammatory activation patterns observed in conventional monocultures and cocultures. However, in contrast to conventional cultures, glial cells cultured in a microfluidic chip revealed enhanced glial crosstalk upon TNF-α/IL-1β stimulation in the microenvironment.

### Glial crosstalk elevates complement component C3 levels in cocultures

One mechanism by which microglia and astrocytes communicate and potentially participate in the progression of neuroinflammatory pathologies is through complement system activation^9,34^. Since we observed reciprocal modulation of inflammatory responses in glial cells, we investigated complement C3 expression in our cultures. Immunocytochemical staining of C3d, a downstream product of C3, revealed high expression in MG monocultures at the basal level and after LPS and TNF-α/IL-1β stimulation (Fig. 7A). AS monocultures and MG/AS cocultures, respectively, demonstrated modest/non-detectable expression of C3d under control conditions. While LPS stimulation had no observable effects, TNF-α/IL-1β stimulation demonstrated strong responses in AS monocultures and MG/AS cocultures, suggesting high expression of C3 in inflammatory astrocytes.

**Figure 7.**
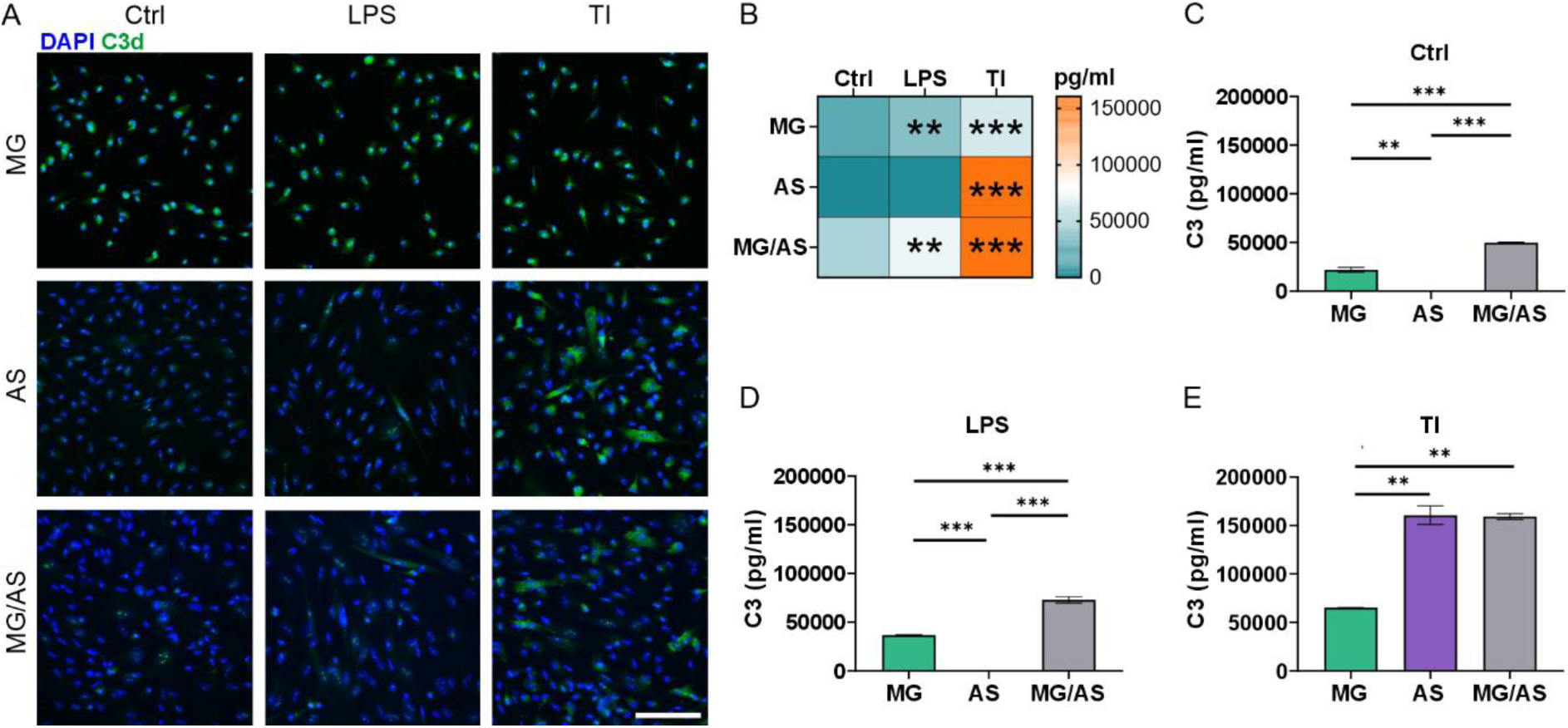
Secretion of complement component 3 (C3) in glial cultures after 24 hours of inflammatory stimulation. **A** Representative images of immunocytochemical staining for C3d in glial monocultures and cocultures. Scale bar, 100 µm. **B** Heatmap of C3 secretion in glial monocultures and cocultures. The data are presented as the means. Asterisks indicate statistical significance in comparison with the control condition. **C** Secretion of C3 in glial cultures under control conditions. **D** Secretion of C3 in glial cultures after LPS stimulation. **E** Secretion of C3 in glial cultures after TNF-α/IL-1β (TI) stimulation. The data are presented as the means ± SDs. n = 2 for all data with 2 technical replicates. *p< 0.05, **p< 0.01, ***p< 0.001; one-way ANOVA with Tukey’s post hoc comparison.

Analysis of C3 secretion levels in the cultures provided further support for the observed staining results (Fig. 7B–E). In the basal state, only the MG monocultures and MG/AS cocultures secreted C3, and this level of secretion increased with both LPS and TNF-α/IL-1β stimulation (Fig. 7B). Interestingly, a greater level of C3 was observed in MG/AS cocultures than in MG monocultures both under control conditions and after LPS stimulation (Fig. 7C and D), indicating the amplification of C3 through microglial crosstalk with astrocytes. As observed with C3 staining, AS monocultures and MG/AS cocultures presented highly elevated secretion of C3 in response to TNF-α/IL-1β stimulation (Fig. 7E). Overall, both MG and AS monocultures, as well as MG/AS cocultures, demonstrated the highest levels of C3 secretion with TNF-α/IL-1β stimulation (Fig. 7C–E), suggesting that TNF-α and/or IL-1β are potent activators of the complement system in both glial cell types. In summary, these results demonstrated complement activation in our glial cultures after inflammatory stimulation as well as the modulation of C3 expression via glial crosstalk.

## Discussion

Increasing evidence indicates a central role of microglia- and astrocyte-driven neuroinflammatory processes in the progression of various neurodegenerative pathologies^21,35^. However, an insufficient understanding of how microglia and astrocytes communicate and mediate inflammatory responses highlights the necessity of developing *in vitro* models that are more physiologically relevant to humans. To address these issues, we established *in vitro* glial coculture models to investigate the function and crosstalk of iPSC-derived microglia and astrocytes under inflammatory conditions. We cocultured glial cells in both conventional culture dishes and a novel microfluidic chip platform and stimulated the cells with LPS, TNF-α/IL-1β, or IFN-γ. LPS is a widely used stimulant of microglial activation and triggers the production and secretion of proinflammatory factors such as TNF-α and IL-1β^10,11,36^. The prototypic inflammatory cytokines TNF-α and IL-1β play significant roles in the activation of proinflammatory astrocytes^26,37,38^. IFN-γ, in turn, is a proinflammatory cytokine with a role in glial activation during neuroinflammatory conditions, including in MS^39,40^. Here, microglia and astrocytes demonstrated strong cell type-specific responses to LPS and TNF-α/IL-1β stimulation that were modulated under coculture conditions, indicating reciprocal signaling between glial cells. In contrast to conventional cultures, the microfluidic chip platform provides a more functional approach for investigating glial crosstalk, enabling the establishment of separate inflammatory microenvironments with spontaneous microglial migration.

The inflammatory responses of microglia and astrocytes have been studied extensively in monocultures via the use of various inflammatory stimulants as well as conditioned media^9–11,26^. Although these studies provide information from unidirectional aspects of inflammatory glial crosstalk, accumulating evidence from coculture studies indicates the importance of direct cell–cell contacts in modulating the inflammatory functions of glial cells^34,35,41^. To understand how microglia and astrocytes modulate the inflammatory responses of the other, we first measured the secretion of inflammatory mediators from the culture media of both conventional monocultures and glial cocultures. In line with previous studies, microglia and astrocyte monocultures responded to LPS and TNF-α/IL-1β stimulation, respectively, by secretion of a wide range of chemokines and cytokines^10,24,36,37^. The observed responses demonstrated cell type specificity, with only minor effects of TNF-α/IL-1β stimulation on microglia and LPS stimulation on astrocytes. While both stimuli elicited broad responses in glial cocultures, alterations in secretion levels suggested the modulation of inflammatory responses via reciprocal signaling. Compared with MG monocultures, LPS stimulation of cocultures induced lower secretion of several inflammatory mediators, indicating dampening of inflammatory responses. This could be due in part to the secretion of the anti-inflammatory cytokine IL-10 by LPS-stimulated microglia, which has been reported to evoke anti-inflammatory signaling between microglia and astrocytes to reduce inflammatory activation^42^. An increase in IL-10 secretion was also observed in the TNF-α/IL-1β-stimulated cocultures, which was indicative of microglial activation through inflammatory astrocytes. Conversely, LPS stimulation induced the secretion of CXCL10 and GM-CSF in cocultures, which was observed predominantly in TNF-α/IL-1β-stimulated astrocyte monocultures, indicating the activation of astrocytes through inflammatory microglia. Previous studies have reported that reactive astrocytes secrete CXCL10 and GM-CSF in chronic CNS neuroinflammation, potentially enhancing the activation of astrocytes as well as the activation of microglia^43–45^. Therefore, these results confirm the intricate reciprocal modulation of inflammatory responses between cocultured microglia and astrocytes.

Since conventional cultures cannot recapitulate the complexity of glial functions within the CNS environment, we next cocultured microglia and astrocytes in a compartmentalized microfluidic chip platform. This approach provided increased control over the conventional culture setup and enabled investigation of glial crosstalk at a more functional level. Microglia and astrocytes were cultured in dedicated compartments in cell type-specific culture media. Chip design allowed the migration of microglia toward astrocytes, enabling the spontaneous formation of glial cocultures. Furthermore, fluidic isolation on the chip platform supported the establishment of separate inflammatory microenvironments, as demonstrated by the selective stimulation of the MG/AS compartment with TNF-α/IL-1β. The stimulation induced high secretion of IL-6, which did not diffuse to the outer compartments over a 7-day extended stimulation period. We further demonstrated the functional potential of the coculture platform with selective IFN-γ stimulation, which has previously been reported to suppress microglial phagocytosis^11^ and markedly reduced phagocytic function of migrated microglia without affecting microglia in unstimulated outer compartments. When treated with inflammatory stimulants, similar to conventional cultures, chip cultures reproduced comparable patterns of inflammatory activation, confirming the applicability of the platform for investigating glial crosstalk. Furthermore, unique responses in chip cocultures suggested an enhanced setup for investigating glial interactions. Although microfluidic chip platforms have not yet been widely utilized for investigating glial functions, their potential for establishing of more physiologically relevant CNS *in vitro* models as well as for disease modeling has been demonstrated in previous studies by us and others^21,24,46^.

As highly motile cells, microglia respond to neuroinflammatory events by migrating toward sites of inflammatory insult and pathological stimuli. Therefore, we investigated how the utilized inflammatory stimulants affected microglial migration rates in our chip coculture platform. The inflammatory stimulants LPS, TNF-α/IL-1β, and IFN-γ had no significant effects on the microglial migration rate regardless of whether the microglia were cultured alone or cocultured with astrocytes. Previous studies utilizing LPS as a stimulant have demonstrated conflicting effects on microglial migration, with some reporting increased^47,48^ and others decreased chemokinesis and chemotaxis^49–51^. Combined stimulation with TNF-α and IFN-γ, in turn, has been reported to decrease microglial migration^52^. However, these studies were conducted utilizing rodent microglia and simple scratch-wound or transwell migration assays involving only one cell type. Interestingly, the presence of astrocytes in our chip cocultures seemed to increase the variability in microglial migration, which may have affected the significance of the results. One well-recognized stimulant of microglial migration is extracellular nucleotide adenosine triphosphate (ATP), which participates in microglial recruitment toward CNS injuries^53–56^. Since ATP and the downstream product ADP have been shown to increase both the chemokinesis and chemotaxis of microglia *in vitro*^10,11,33,57^, we utilized ADP stimulation as a positive control in our fluidically isolated chip cocultures. Accordingly, our results demonstrated that the migration of microglia increased after ADP stimulation, confirming the potential of our coculture platform for migration studies.

Activation of the complement system mediates crosstalk between microglia and astrocytes during neuroinflammatory responses^9,34^ and has also been demonstrated in several neuroinflammatory diseases, including AD^58^ and MS^59,60^. In this study, we showed that microglia, but not astrocytes, expressed C3 in the basal state. However, following inflammatory activation, astrocytes expressed C3 more prominently than microglia. Consistent with our results, previous publications have shown that the secretion of C3 is characteristic of reactive neuroinflammatory astrocytes *in vitro*^9,24^. Here, coculturing microglia and astrocytes elicited a significant increase in C3 secretion in comparison to monocultures in both the basal state and after LPS stimulation, demonstrating aggravation via reciprocal signaling between microglia and astrocytes. Accordingly, C3 secretion was observed to be mediated via glial crosstalk in neuronal tricultures in a recent publication by Guttikonda and colleagues^34^. Our results therefore support the role of C3 in inflammatory glial crosstalk, as C3 was upregulated in both microglia and astrocytes upon inflammatory activation and further potentiated in glial cocultures.

## Conclusions

Taken together, our results provide valuable information on the inflammatory molecular conversation between microglia and astrocytes, revealing cell type-specific inflammatory responses as well as reciprocal signaling through secreted chemokines and cytokines in glial cocultures. By coculturing microglia and astrocytes in our compartmentalized microfluidic coculture platform, we were able to explore inflammatory glial interactions at a more functional level within distinct, interconnected inflammatory microenvironments. Microglia efficiently migrated toward astrocytes, forming spontaneous glial cocultures and increasing their migration in response to the chemoattractant ADP. Moreover, the microfluidic coculture platform successfully recapitulated inflammatory activation and crosstalk between microglia and astrocytes, revealing unique inflammatory responses within glial cocultures. Our results further demonstrated elevated secretion of C3 within glial cocultures upon inflammatory activation, emphasizing the role of C3 in inflammatory glial crosstalk. Our *in vitro* coculture platform may be a valuable tool for elucidating the mechanisms of glial function and crosstalk in neuroinflammation.

## Acknowledgements

The authors acknowledge Biocenter Finland (BF), the Tampere Facility of iPS Cells, and the Tampere Imaging Facility (TIF) for their service (Faculty of Medicine and Health Technology, Tampere University). We thank Katriina Aalto-Setälä (Heart Group, Tampere University) for providing the healthy control iPSC line. We also thank Eija Nieminen, Outi Heikkilä and Hanna Mäkelä for their technical assistance with cell maintenance and analyses. Furthermore, we thank Begum Gokce for her technical assistance with astrocyte differentiation.

## Author contributions

I.T., T.H. and S.H. designed the study; P.K. provided the microfluidic chips produced by L.S. and K.T.; J.R., D.V. and A.H. provided astrocytes and assisted with the astrocyte differentiation protocol; H.J. and T.M. provided and assisted with the microglia differentiation protocol; I.T. and K.K. differentiated astrocytes with help from T.H.; J.L., T.H. and S.H. performed the differentiation of microglia; I.T., T.H. and S.H. performed the experiments with help from J.L.; I.T., T.H. and S.H. analyzed the data; I.T., T.H. and S.H. wrote the manuscript; T.M., A.H., P.K., S.N. and S.H. provided resources for the study. All the authors edited and approved the manuscript.

## Funding

This work was supported by the Research Council of Finland (SH 330707, 335937, and 358045, SN 36665 and PK 336785), the Finnish MS Foundation (SH and JL), the Päivikki ja Sakari Sohlberg Foundation (SH), the Finnish Cultural Foundation (SH, TH and IT), the Maud Kuistila Memorial Foundation (JL), the Orion Research Foundation sr (JL and IT), the Tampere Institute of Advanced Study (TH), the Doctoral Programme in Medicine, Biosciences and Biomedical Engineering, Tampere University (JL and IT), and the European Union (HORIZON-MSCA-2022-PF-01, GA-101109010-NEoC awarded to JR).

## Availability of data and materials

All data are available in the main text or the supplementary material.

## Competing interests

The authors report no competing interests.

## Notes

### Competing Interest Statement

The authors have declared no competing interest.

