## Supplementary material for "Human iPSC-based coculture model reveals neuroinflammatory crosstalk between microglia and astrocytes"

#### Supplementary Figures

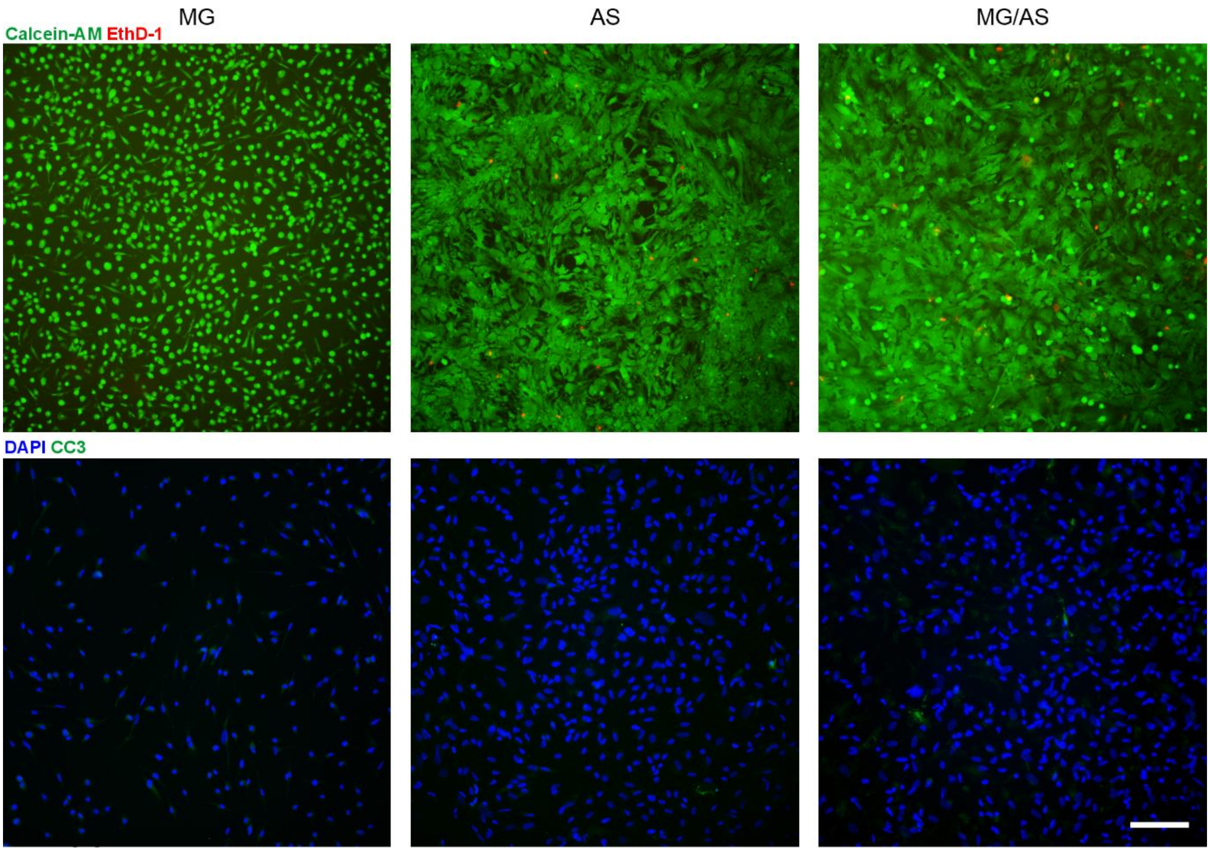


**Supplementary Figure 1. Viability of glial monocultures and cocultures.** Representative immunofluorescence images of cells stained with the viability markers calcein-AM and EthD-1 and the apoptotic marker cleaved caspase-3 (CC3). Scale bar, 100 µm.

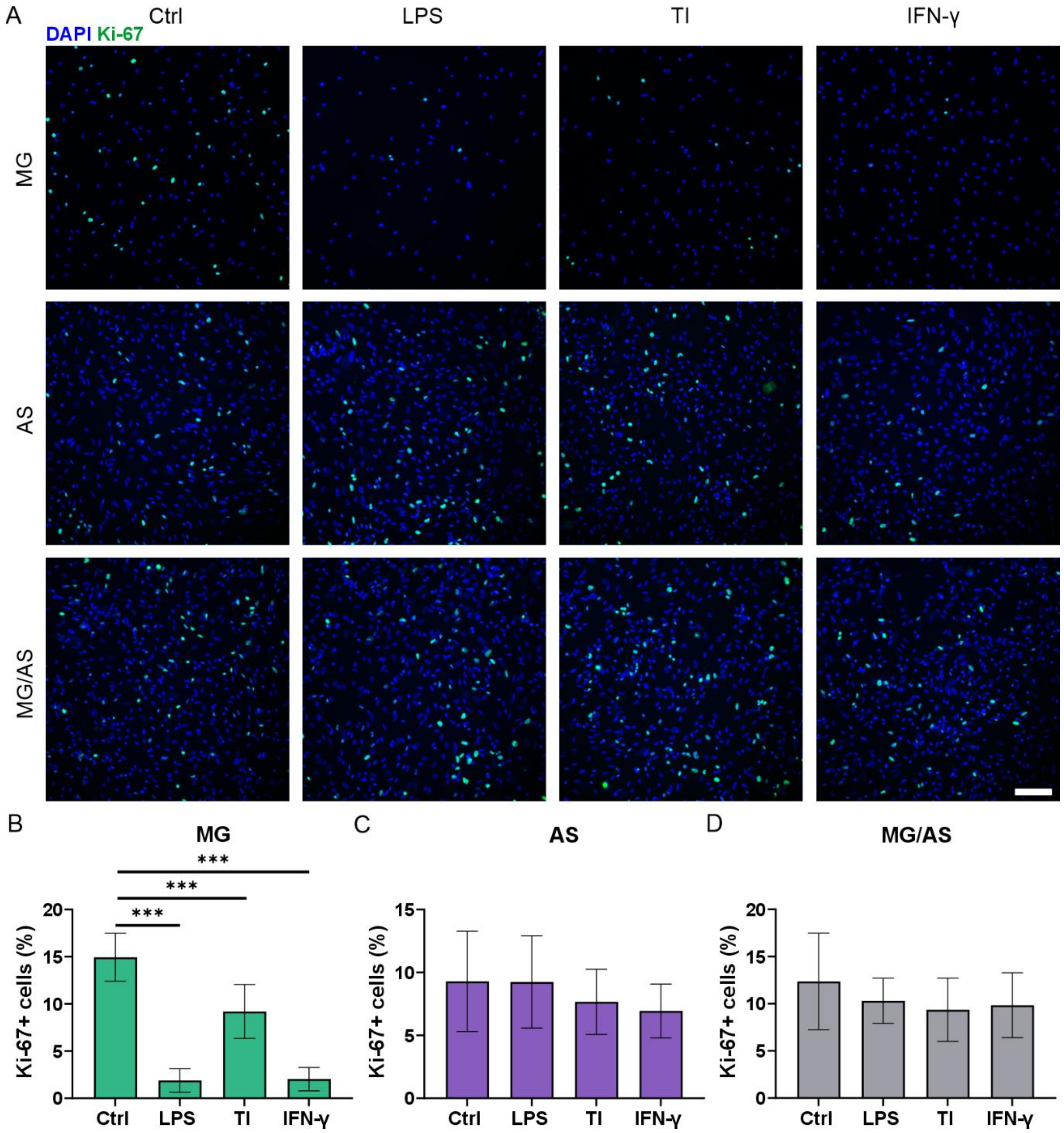


**Supplementary Figure 2. Proliferation of glial cultures after 24 hours of inflammatory stimulation. A** Representative images of immunocytochemical staining for the proliferation marker Ki-67. Scale bar, 100 µm. **B–D** Ki-67-positive (+) cells were quantified from cultures as a percentage (%) of the total cell count. n = 18 images per condition; images from 2 experiments. The data are presented as the means ± SDs. ***$p < 0.001$; one-way ANOVA with Tukey's post hoc comparison.

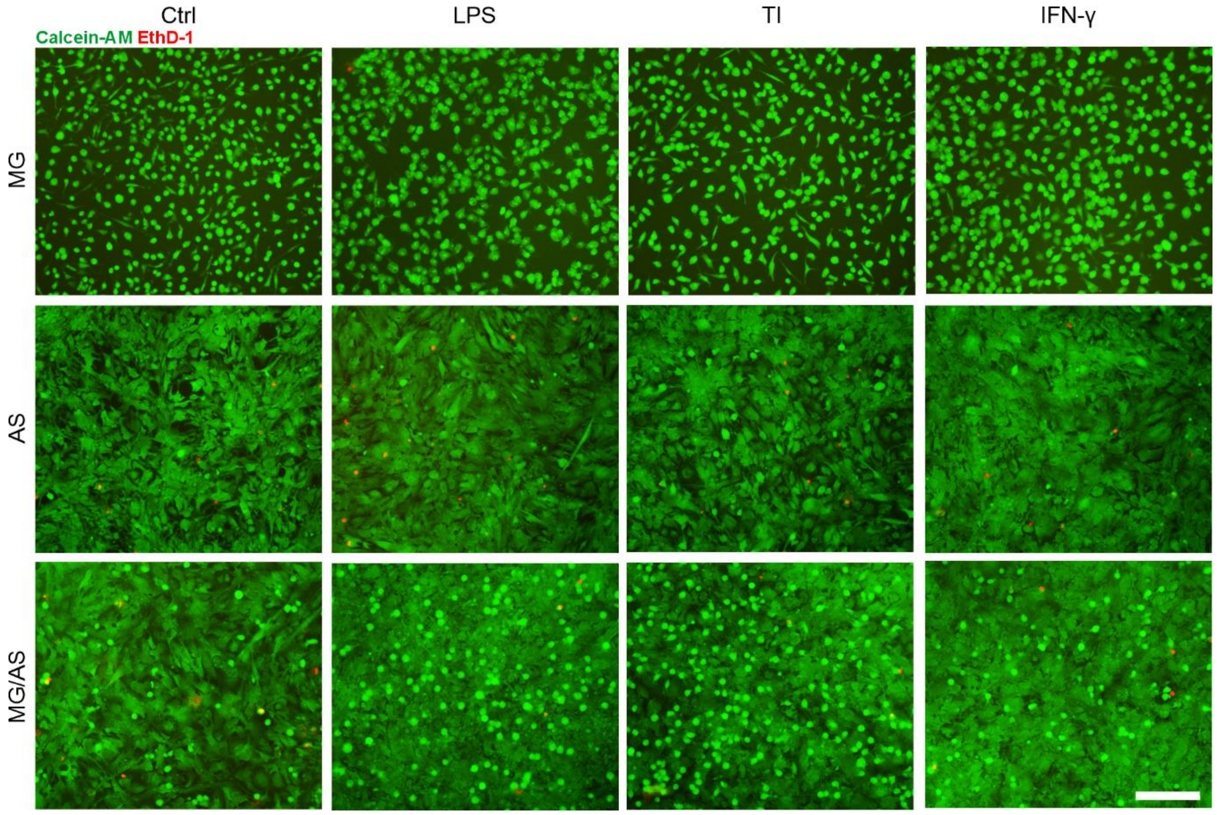


**Supplementary Figure 3. Viability of glial cultures after 24 hours of inflammatory stimulation.** Representative immunofluorescence images of cells stained with the viability markers calcein-AM and EthD-1. Scale bar, 200 μm.

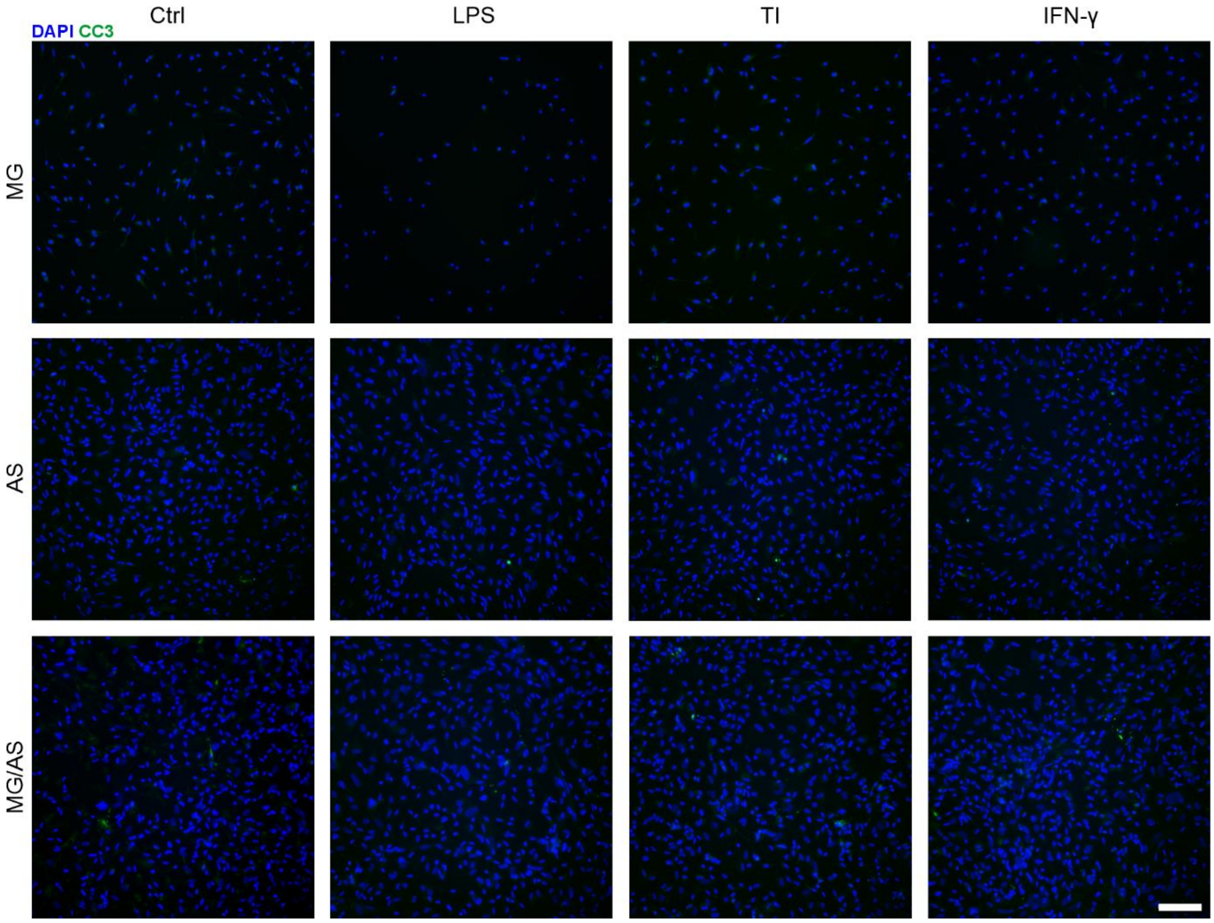


**Supplementary Figure 4. Staining of the apoptotic marker cleaved caspase-3 (CC3) in glial cultures after 24 hours of inflammatory stimulation.** Representative images of immunocytochemical staining for the apoptotic marker CC3. Scale bar, 100 µm.

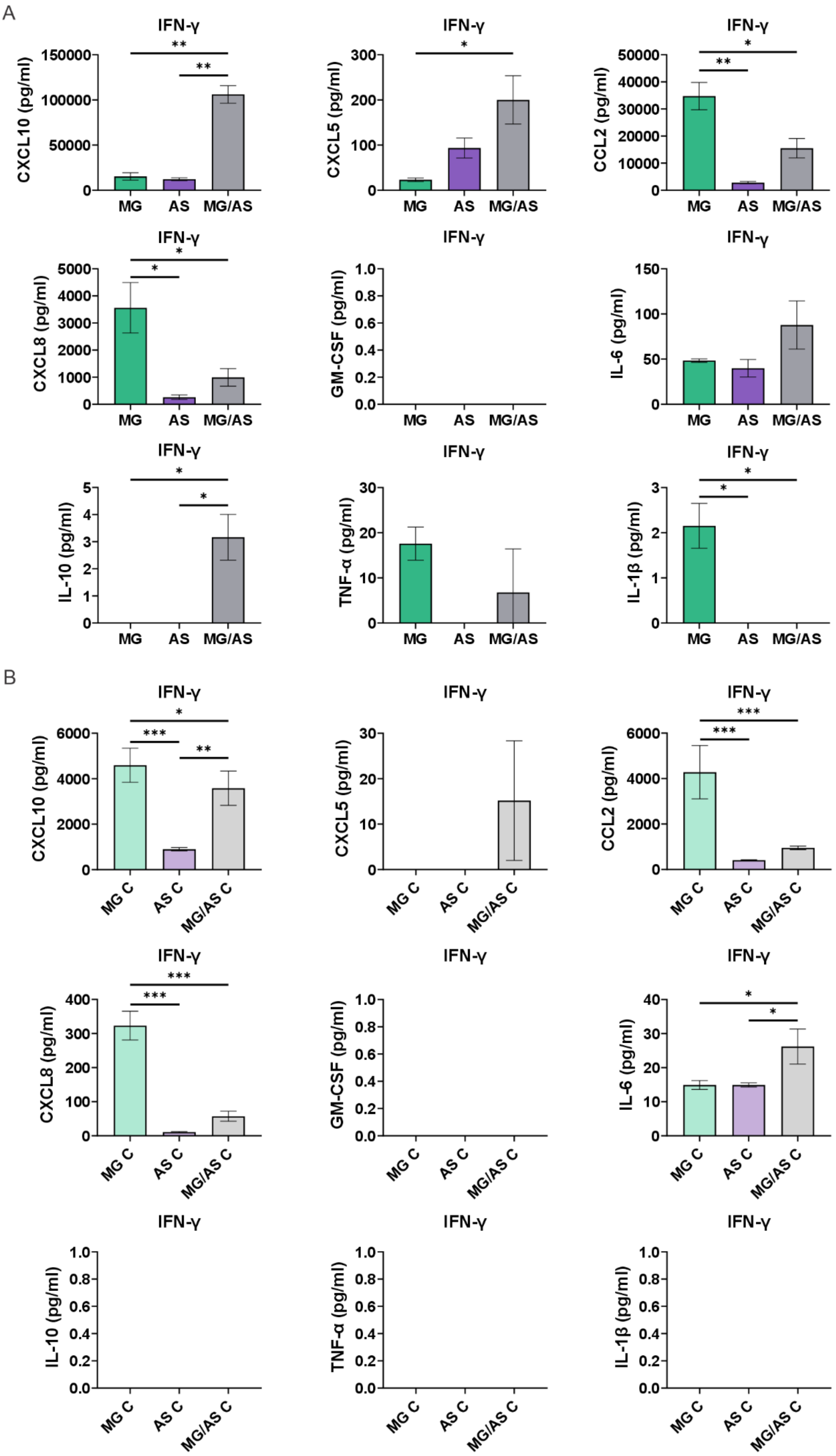
A
IFN-γ
CXCL10 (pg/ml)
150000
100000
50000
0
MG AS MG/AS
**
**
IFN-γ
CXCL5 (pg/ml)
300
200
100
0
MG AS MG/AS
*
IFN-γ
CCL2 (pg/ml)
50000
40000
30000
20000
10000
0
MG AS MG/AS
*
**
IFN-γ
CXCL8 (pg/ml)
5000
4000
3000
2000
1000
0
MG AS MG/AS
*
*
IFN-γ
GM-CSF (pg/ml)
1.0
0.8
0.6
0.4
0.2
0.0
MG AS MG/AS
IFN-γ
IL-6 (pg/ml)
150
100
50
0
MG AS MG/AS
IFN-γ
IL-10 (pg/ml)
5
4
3
2
1
0
MG AS MG/AS
*
*
IFN-γ
TNF-α (pg/ml)
30
20
10
0
MG AS MG/AS
IFN-γ
IL-1β (pg/ml)
3
2
1
0
MG AS MG/AS
*
*
B
IFN-γ
CXCL10 (pg/ml)
6000
4000
2000
0
MG C AS C MG/AS C
*
***
**
IFN-γ
CXCL5 (pg/ml)
30
20
10
0
MG C AS C MG/AS C
IFN-γ
CCL2 (pg/ml)
6000
4000
2000
0
MG C AS C MG/AS C
***
***
IFN-γ
CXCL8 (pg/ml)
400
300
200
100
0
MG C AS C MG/AS C
***
***
IFN-γ
GM-CSF (pg/ml)
1.0
0.8
0.6
0.4
0.2
0.0
MG C AS C MG/AS C
IFN-γ
IL-6 (pg/ml)
40
30
20
10
0
MG C AS C MG/AS C
*
*
IFN-γ
IL-10 (pg/ml)
1.0
0.8
0.6
0.4
0.2
0.0
MG C AS C MG/AS C
IFN-γ
TNF-α (pg/ml)
1.0
0.8
0.6
0.4
0.2
0.0
MG C AS C MG/AS C
IFN-γ
IL-1β (pg/ml)
1.0
0.8
0.6
0.4
0.2
0.0
MG C AS C MG/AS C

**Supplementary Figure 5. Secretion of inflammatory factors after 24 hours of IFN-γ stimulation in MG and AS monocultures and MG/AS cocultures. A** Comparison of the secretion levels of cytokines and chemokines between different conventional cultures stimulated with IFN-γ. n = 2, with 2 technical replicates; the data are representative of 2 experiments. The data are presented as the means ± SDs. **B** Comparison of the secretion levels of cytokines and chemokines between different cell compartments in the chip after IFN-γ stimulation. n = 2–3, with 2 technical replicates; the data are representative of 2 experiments. The data are presented as the means ± SDs. *p< 0.05, **p< 0.01, ***p< 0.001; one-way ANOVA with Tukey's post hoc comparison.

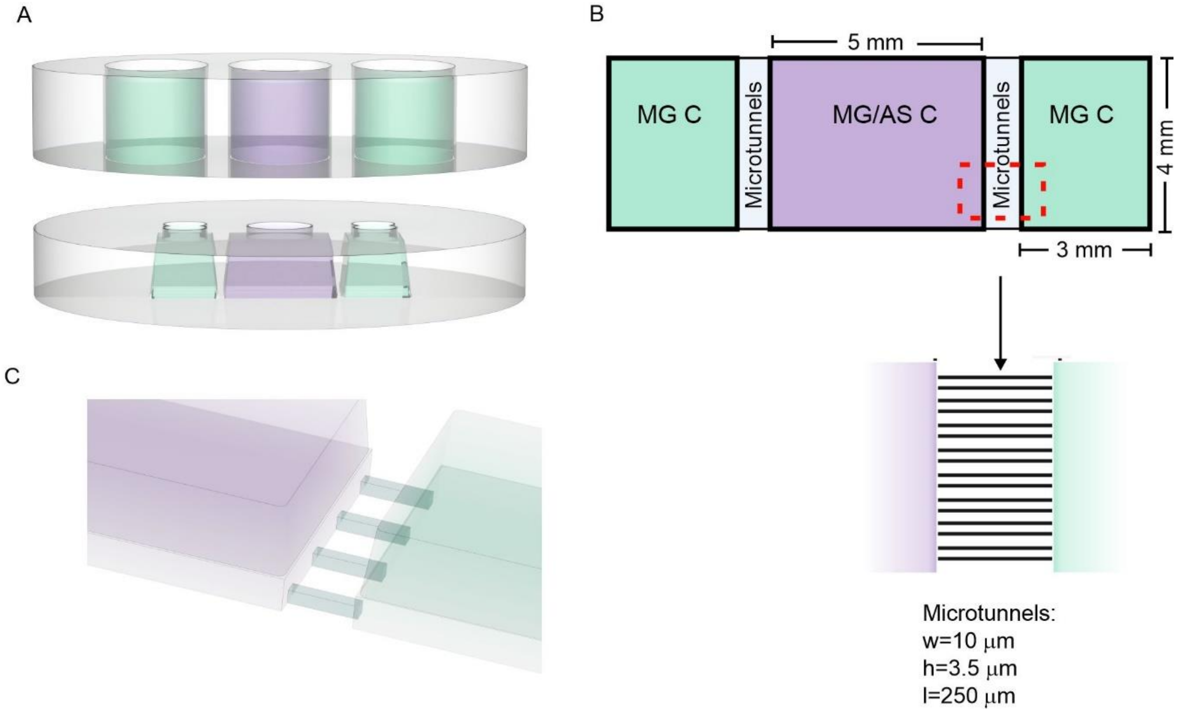


**Supplementary Figure 6. Design of the microfluidic platform. A** The microfluidic platform is composed of two separate PDMS parts: a medi reservoir (upper) and a cell culture part (lower) containing three cell compartments. **B** The microglia/astrocyte compartment (MG/AS C) is interconnected with 40 microtunnels to two microglia compartments (MG C). The dimensions of the cell compartments and microtunnels (l=length; h=height, w=width) are described in the image. **C** Close-up image showing microtunnels connecting to the cell compartments.

### Supplementary Tables

**Supplementary Table 1.** Primary and secondary antibodies used for the immunocytochemical stainings.

| Primary antibody | Animal | Product number | Producer | Dilution 2D | Dilution chips |
|---|---|---|---|---|---|
| Iba1 | Rabbit | 019-19741 | FujiFilm Wako | 1:500 | 1:200 |
| TMEM119 | Rabbit | ab185333 | Sigma-Aldrich | 1:100 | - |
| P2RY12 | Rabbit | HPA014518 | Sigma-Aldrich | 1:125 | - |
| CD44 | Rabbit | ab157107 | Abcam | 1:500 | - |
| S100β | Mouse | S2532 | Sigma-Aldrich | 1:500 | 1:200 |
| GFAP | Chicken | ab4674 | Abcam | 1:4000 | - |
| Ki-67 | Rabbit | AB9260 | Millipore | 1:800 | - |
| Cleaved Caspase-3 | Rabbit | 9664 | Cell signaling Technology | 1:400 | - |
| C3d complement | Rabbit | A0063 | Agilent Technologies | 1:2000 | - |
| **Secondary antibody** | **Animal** | **Product number** | **Producer** | **Dilution 2D** | **Dilution chips** |
| Alexa Fluor 488 donkey anti-rabbit IgG (H+L) | Donkey | A21206 | Thermo Fisher Scientific | 1:400 | 1:200 |
| Alexa Fluor 488 donkey anti-mouse IgG | Donkey | A21202 | Thermo Fisher Scientific | 1:400 | 1:200 |
| Alexa Fluor 568 donkey anti-mouse IgG | Donkey | A10037 | Thermo Fisher Scientific | 1:400 | 1:200 |
| Alexa Fluor 568 donkey anti-rabbit IgG | Donkey | A10042 | Thermo Fisher Scientific | 1:400 | 1:200 |
| Alexa fluor 647 goat anti-chicken IgY | Goat | A21449 | Thermo Fisher Scientific | 1:200 | - |
